# Stress History Modulates CRF Neurons to Establish Resilience

**DOI:** 10.1101/2022.08.31.505596

**Authors:** Sherod E Haynes, Anthony Lacagnina, Hyun Seong Seo, Muhammad Afzal, Carole Morel, Aurelie Menigoz, Kanaka Rajan, Roger L Clem, Barbara Juarez, Helen S Mayberg, Donald G. Rainnie, Larry J. Young, Ming-Hu Han

## Abstract

**Introduction:** Cumulative stress is a major risk factor for developing major depressive disorder (MDD), yet not everyone experiencing chronic stress develops MDD. In those who do not, it is unclear at what point, or by what mechanism, a trajectory of stable resiliency emerges.

**Methods:** Utilizing a 10-day repeated social defeat stress model (RSDS) for MDD, we observed that a critical period between 7 and 10 daily defeats marks the phenotypical divergence of resilient from susceptible mice. Using cell-type selective electrophysiology, chemogenetics, optogenetics, fiber photometry and RNA quantification was employed to investigate the nature of stress effects on neuroadaptation in the oval nucleus of the bed nucleus of the stria terminalis (BNSTov) required to determine resilience.

**Results:** In response to ongoing stress, corticotropin-releasing factor (CRF^+^, but not CRF^-^) neurons of the (BNSTov) displayed a sustained increased firing rate in resilient, but not susceptible mice. This neurophysiological adaptation was self-sustaining, but only after 7 critical stress exposures, indicating that the process of developing resilience is dependent on stress history.

**Conclusion:** Our study reveals a novel process by which individuals might persist in the face of adversity by way of stress-provoked activation, not inhibition of a key CRF limbic region that establishes a pathway to resilience.

## Introduction

Major depressive disorder (MDD) is a crippling heterogeneous neuropsychiatric condition with high morbidity and lifetime prevalence.^1,2^ Major life stressors are key precipitants in the onset of an episode^3,4^. While most people will report at least one major life stressor at some point in their lives, not everyone will go on to develop MDD. It is also known that depression can emerge after significant repeated stressor exposure, suggesting that cumulative exposure to stress is associated with increased depression risk in vulnerable individuals^5–8^. Cumulative stress causes numerous psychological insults that predispose neuropsychiatric conditions^5,9,10^. Many studies have explored the individual differences in stress susceptibility or resiliency, but the process critical to driving the divergence in phenotypes remain elusive. With the odds of treatment failure increasing with subsequent depressive episodes,^2,12,13^ and increased depression risk of cumulative stress, a potentially more effective strategy would be to target the mechanisms mediating resilience, enhancing the ability to cope with cumulative stress.

Resiliency is defined as the "process of adapting well in the face of adversity… [to] threats, or significant sources of stress.”^14–16^ Resiliency is unlikely to emerge from the absence of a pathological stress response but instead is an active process involving a myriad of molecular, cellular and circuit-level changes in the brain. Repeated social defeat stress (RSDS) induces robust depression-like behavioral phenotypes in roughly 2/3 of mice^6,10^. The standard 10-day RSDS protocol has been employed widely to identify and study neurobiological features of susceptible and resilient subpopulations^6,7,17–23^. Studies on stress vulnerability often test hypotheses after establishing vulnerable subgroups, making the processes that led to such phenotypic diversification unclear.

The Bed Nucleus of the Stria Terminalis (BNST) is an important node for integrating sensory cues, interoception, cognition, and motivational states to enact adaptive responses^24,25^. The BNST is well-positioned in the social salience network to integrate external cues with internal states to influence the outcome and context of social interactions^24–27^. The oval nucleus of the BNST (BNSTov) is a stress-sensitive subregion that is a key candidate region for encoding stress modulation of social behavior through its projections to areas such as the ventral tegmental area (VTA)^28–31^ and dorsal raphe^32,33^. Corticotrophin-releasing factor (CRF) neurons of the BNSTov (BNSTov^CRF^) are a significant output of this region and can influence affective states^34–37^. Since BNSTov^CRF^ neurons are thought to promote arousal and adaptive responding according to stress-related changes in internal state^24,25,27,38–44^, we hypothesized they might play a role in establishing resiliency to repeated social stress.

Using the 10-day RSDS paradigm, we used cell-type selective *ex vivo* electrophysiology, chemogenetics, optogenetics, and *in vivo* fiber photometry to interrogate the BNSTov^CRF^ system to explore its role in the divergence of susceptible and resilient phenotypes. We observed that BNSTov^CRF^ neurons encode repetitive social stress and undergo adaptation that coincides with the divergence of resilient and susceptible phenotypes. Unexpectedly, we observed that resiliency entails cumulative stress-history dependent neuroadaptive changes, whereas susceptibility emerges in the absence of similar adaptations. Activation of BNSTov^CRF^ neurons using Designer-Receptors-Exclusively-Activated-by-Designer-Drugs (DREADDs) during social defeat led to a resilient phenotype while inhibition caused susceptibility. Finally, using RNAscope and optogenetics, we provide intriguing evidence for a potential role of *Crfr1* in BNSTov^CRF^ neurons in mediating resilience.

## Methods & Materials

### Mice

The study used wild-type, Crf-ires-Cre (Jackson labs: 011087), ai14 (Cre-responsive tdTomato reporter mouse; Jackson labs: 007915), ai32 (Cre-responsive channelrhodopsin-2/fused with eYFP; Jackson labs: 012569) mice on C57BL/6J background that were bred at Icahn School of Medicine at Mount Sinai and used were between 6-7 weeks at the start of experimental manipulations. All experiments were approved by the Institutional Animal Care and Use Committee and comply with institutional guidelines for the Animal Care and Use Committee set forth by Icahn School of Medicine at Mount Sinai.

### Repeated social defeat stress paradigm

The repeated social defeat stress paradigm was performed according to published protocols^6,21,23,82,90–92^. Briefly, CD1 aggressors were singly housed in cages (26.7 cm width x 48.3 cm depth x 15.2 cm height; Allentown Inc) at least 24 hours before the start of the experiment on one side with a clear perforated Plexiglas divider. Social stress consists of physical and sensory stress components.

### Viral constructs

For DREADD experiments in CRF neuronal populations, CRF-ires-Cre animals were injected with AAV5-hSyn-DIO-hM4D(Gi)-mCherry (Addgene: 44362-AAV5) (≥7x10^12), AAV5-hSyn-DIO-hM3D(Gq)-mCherry (Addgene: 44361-AAV5) (≥7x10^12), and AAV5-hSyn-DIO-mCherry (Addgene: 50459). For fiber photometric recordings, CRF-ires-Cre animals were injected with AAV9-syn-FLEX-jGCaMP7f-WPRE (addgene: 104492-AAV9) ( >1x10^12). All viruses were purchased from Addgene.

### CNO-drinking water construct

Clozapine-N-Oxide (CNO) was obtained from Hello Bio (catalog no HB6149). The dry chemical was dissolved in drinking water obtained from the vivarium and diluted such that each mouse received 5 mg/kg/day based on previous studies. CNO was made fresh each day for the three days it was administered. CNO solutions were protected from light throughout the experimental procedure. On average, mice consumed ∼4-5 mL of water per day. Water bottles and mice were weighed daily.

Please see supplementary methods for further details.

## Results

### Susceptible vs. resilience phenotypes emerge between 7 and 10 daily episodes of social defeat stress

RSDS produces divergent and enduring resilient/susceptible phenotypes following 10 episodes of social defeat stress^6,21^ and enables the investigation of the consequences of stress accumulation^5,7^. To examine the temporal divergences of resilient and susceptible phenotypes in mice, a modified RSDS stress protocol was employed in which mice were subjected to discrete numbers of social defeat episodes (SDEs) interspersed with social interaction (SI), and sucrose preference (SP) tests administered after 1, 4, 7, and 10 SDEs (Fig. 1A-D). Social Interaction (SI) Ratio is a behavioral score of the SI test and measures the time spent in the area proximal to the enclosure of a novel social target (social interaction zone). To generate the SI ratio, interaction zone time is calculated both in the absence and presence of the social target, such that SI ratio ≧ 1 is resilient and <1 is susceptible^21^. Surprisingly, we found that the susceptible phenotype emerged discretely between 7 and 10 SDEs (Fig. 1E, Supplemental Data Fig 1A) (SI ratio: 2.062 +/- 0.226 after 7 SDEs to 0.559 +/- 0.093, p<0.0001, n = 11 for susceptible and 1.967 +/- 0.188 (7 SDEs) to 1.554 +/- 0.163 (10 SDEs), p=0.2929, n=11 for resilient). There were no significant differences in SI ratios after 7 SDEs between mice that went on to become susceptible vs resilient after 10 SDEs (mean: 2.062 +/- 0.226 vs 1.967 +/- 0.188, two-tailed t-test, p=0.7483, n= 11/group) (Supplemental Data Fig. 1B-C). This effect was not due to repeated SI tests (Supplemental Data Fig. 2D-F). Resilient mice had an indistinguishable SI ratio from all mice after 7 SDEs, while susceptible (10 SDEs) mice had an SI ratio significantly lower than both groups (Fig. 1F, G). In the sucrose preference (SP) test, resilient mice showed a similar preference for sucrose to all 7 SDE and stress-naïve control mice. Previous studies reported that RSDS produces susceptible and resilient phenotypes after 10 SDEs in a bimodal distribution^21^. Here, we observed a unimodal distribution of social interaction toward a novel conspecific in 7 SDE mice but a bimodal distribution in 10 SDE mice, consistent with the emergence of distinct resilience and susceptible phenotypes (Kolmogorov-Smirnov, p<0.0001; Fig. 1 (H,I,J). The time spent interacting with a novel conspecific was significantly less in susceptible 10 SDE mice compared to the 7th SDE and 10th SDE resilient mice, which were indistinguishable (Fig.1K). The phenotypic divergence was dependent upon the number of SDEs rather than the passage of time and dependent upon the anterior dorsal BNST(Supplemental Data Fig. 1G, 2A-E).

**Figure 1.**
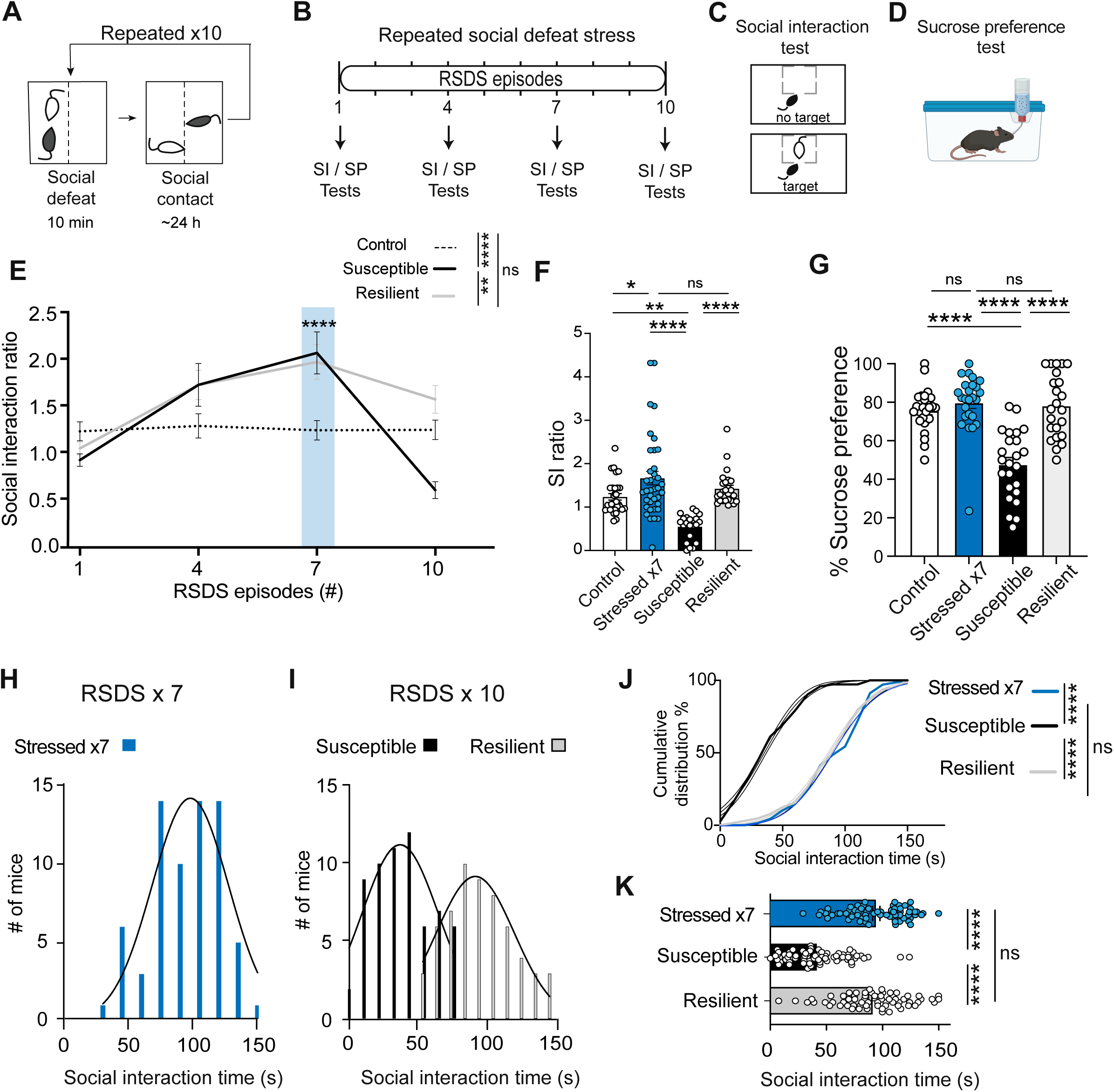
Susceptible and resilient subgroups emerge between 7 and 10 daily episodes of social defeat stress. **A.** Experimental design for repeated social defeat stress (RSDS). **B.** Experimental timeline of social defeat stress and behavioral tests: social interaction (SI) and sucrose preference (SP). **C.** Social interaction test schema involving target and no target trials. **D.** Sucrose preference test schematic. **E.** Effect of cumulative social defeat stress on social interaction measured as SI ratios, after 1, 4, 7, and 10 social defeat episodes (SDEs) (n=11-18 mice/group), two-way ANOVA interaction F_(6,134)_=5.783, ****P<0.0001, row factor F_(3,134)_=12.11, ****P<0.0001, F_(2,134)_=4.086, column factor *P<0.05, Tukey’s post-hoc test susceptible vs resilient ****P<0.0001, susceptible vs control **P<0.01, resilient vs control P=0.2751. SI test susceptible (SI test stressed x7 vs stressed x10) ****P<0.0001. **F.** Aggregated data on social interaction test across experiments. One-way ANOVA treatment F_(3,109)_=14.61, ****P<0.0001. Tukey’s post-hoc test control vs stressed x7 *P<0.05, control vs susceptible **P<0.01, stressed x7 vs susceptible ****P<0.0001, susceptible vs resilient ****P<0.0001 (n=25-37 mice). **G.** Sucrose preference test. One-way ANOVA treatment F_(3,93)_=24.06, ****P<0.0001). Tukey’s post-hoc test control vs stressed x7 P=0.77, control vs susceptible ****P<0.0001, control vs resilient P=0.9387, stressed x7 vs susceptible ****P<0.0001, stressed x7 vs resilient P=0.984, susceptible vs resilient ****P<0.0001 (n=23-25 mice). **H.** Distribution of stressed mice who underwent 7 SDEs. **I.** Distribution of mice who underwent 10 SDEs and sorted into susceptible and resilient mice**. J.** Cumulative distribution of all stressed mice. Kolmogorov-Smirnov (distance) 0.2054 Susceptible vs Resilient, ****P<0.0001. **K.** Time spent interaction socially with a novel conspecific. One-Way ANOVA F_(2,201)_=76.63, ****P<0.0001. Tukey’s post-hoc susceptible vs resilient ***P<0.001, susceptible vs stressed x7 ****P<0.0001, stressed vs resilient P=0.4984 (n=72 susceptible, 64 resilient mice). All data represent means ± SEM. *P < 0.05, **P < 0.01, ***P < 0.001, ****P<0.0001, ns = not significant.

### Divergence in BNSTov^CRF^ neuronal firing rates tracks the emergence of resilient and susceptible phenotypes

CRF neurons of the BNST are a major output source of the oval BNST that are sensitive to chronic stressors^27,45–47^. Thus, we hypothesized that chronic stress would alter BNSTov^CRF^ neuronal activity, coinciding with the divergence in resilient and susceptible behavioral phenotypes. To test this hypothesis, *Crf-*ires*-Cre;*ai14 (tdTomato) mice^30,48–51^ were subjected to either 7 or 10 daily SDEs, and cell-attached *ex-vivo* electrophysiological recordings were conducted in the BNSTov (Fig. 2A,B). CRF^+^, but not CRF^-^, BNSTov neurons of 7 SDE mice had significantly increased firing rates compared to stress-naïve control mice. By contrast, CRF^+^ firing in susceptible 10 SDE mice was indistinguishable from that of controls and lower than 7 SDE and resilient mice. CRF^-^ neuronal firing did not differ between groups (Fig. 2C,D). Moreover, there was a strong correlation between firing rate and social interaction ratio in CRF^+^ but not CRF^-^ neurons in 10 SDE mice (CRF^+^, R^2^=0.5725, *p=0.0113; CRF-R^2^=0.06096, p=0.5219, Fig. 2E). CRF neurons display both burst and non-burst firing patterns^40,52,53^ (Fig. 2F). Burst firing patterns were prominent in resilient and 7 SDE mice compared to susceptible and control mice (Fig. 2H). Additionally, percentage spikes within burst were higher in 7 SDE and resilient compared to susceptible 10 SDE mice (***p=0.0004, one-way ANOVA, Fig. 2I), but not susceptible or control mice. The number of spikes per burst and number of bursts per cell was not significantly different amongst the groups (Fig. 2J,K). These data and correlation analysis suggest the possibility that there is a causal link between the neuronal activity of BNSTov^CRF^ neurons and the divergence of behavioral phenotypes.

**Figure 2.**
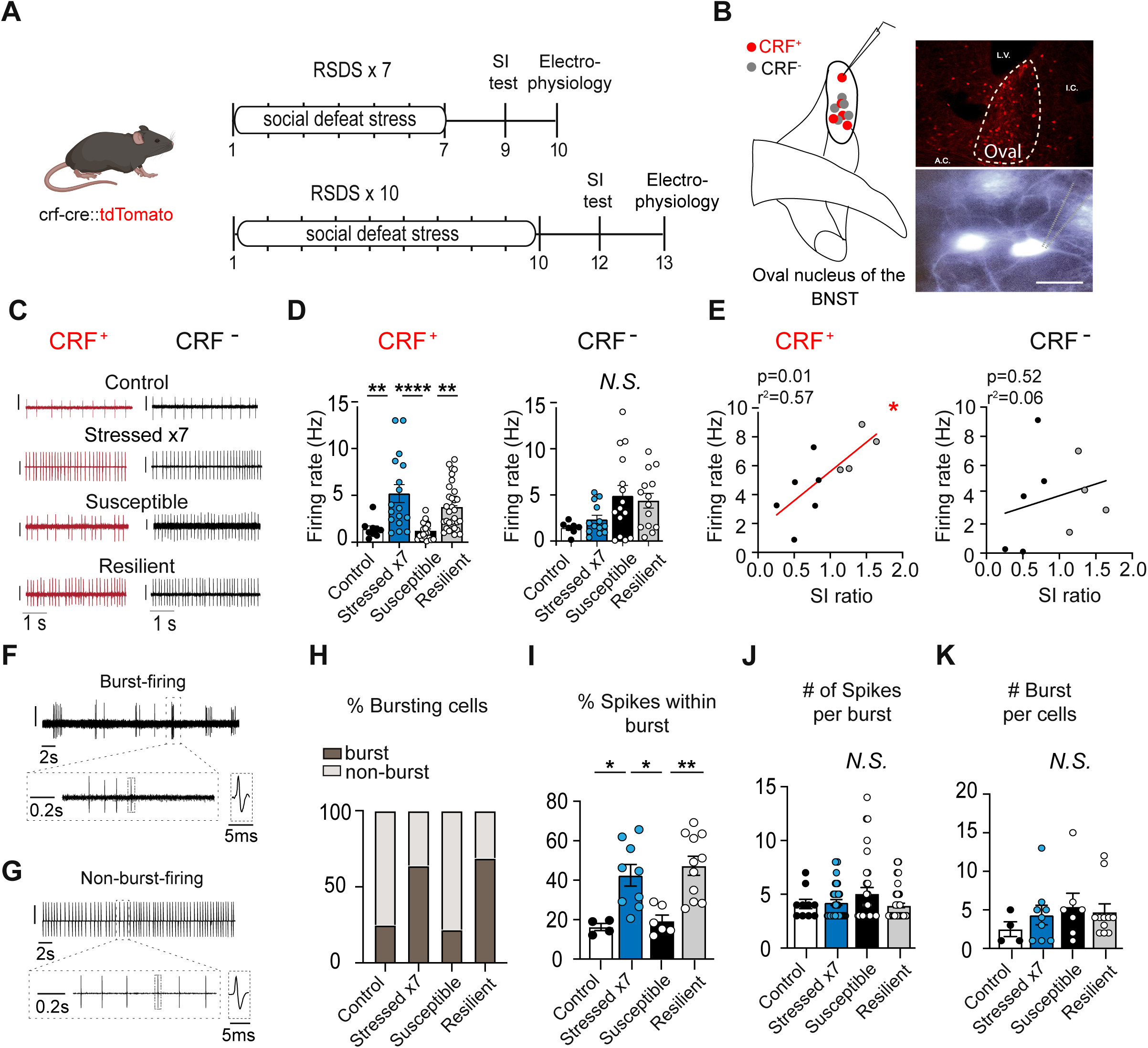
Firing rate alterations in BNSTov^CRF^ neurons occur as adaptation to social stress, persisting in resilient, not susceptible mice. **A.** Mouse genotype and timeline of cell-attached electrophysiology experiments**. B.** Fluorescence-guided cell-attached electrophysiology setup, brain slice of the BNST (CRF cells, tdTomato) and DIC image of CRF neurons, scale bar 0.63 mm. **C.** Representative trace of BNSTov^CRF^ positive and negative neurons of control, stressed (SDEs x 7), susceptible, and resilience mice. **D.** Firing rate of CRF^+^ neurons (n=9-31 cells/group), ****P<0.0001, one-way ANOVA, Tukey’s multiple comparison’s test, control vs stressed (SDEs x 7) **P<0.01, susceptible vs resilient **P<0.01, susceptible vs stressed (SDEs x7), ****P<0.0001. Firing rate of CRF^-^ neurons (n=7-15 cells per 4-6 mice/group), one-way NOVA, F_(3, 45)_=3.113, *P<0.05, Tukey’s multiple comparison’s test, control vs stressed x7 P=0.9192, control vs susceptible P=0.0769, control vs resilient P=0.1742, stressed (SDEs x 7) vs susceptible P=0.1370, stressed x7 vs resilient P=0.3219, susceptible vs resilient P=0.9673. **E.** Correlation of firing rate with social interaction ratio: CRF^+^, R^2^=0.5726, *P<0.05; CRF^-^, R^2^=0.0609, P=0.5219. **F.** Representative sample of burst firing. **G.** Representative sample of tonic firing. **H.** Percentage of bursting cells per animal group. **I.** Percentage of spikes within burst. **J.**Number of spikes per burst. **K.** Number of bursts per cell. (n=10-52 cells per 4-6 mice/group). All data represent means ± SEM. *P<0.05, **P<0.01, ***P<0.001, ****P<0.0001, N.S. = not significant.

### BNSTov^CRF^ neurons bidirectionally modulate the emergence of resiliency

To test the hypothesis that BNSTov^CRF^ neurons regulate and maintain resilience over the last 3 episodes of RSDS, we injected *Crf-*ires-*Cre* mice with AAVs encoding Cre-dependent excitatory (DIO-hM3Dq), inhibitory (DIO-hM4Di) DREADDs, or mCherry construct (control) into the BNSTov and administered CNO via drinking water^54–56^ (Fig. 3A,B). We presaged that this experimental design afforded the opportunity to modulate neurons over a longer time span than intraperitoneal (IP) injections would allow with minimal invasiveness, particularly because it was not clear over when neuroadaptation occurs within the 3-day window. mCherry control mice exhibited both susceptible and resilient phenotypes (SI ratio <1.0 and ≧1.0, respectively) in roughly a 60/40 ratio as expected^6,21^ (Fig. 3C,F). Interestingly, mice injected with inhibitory DIO-hM4Di displayed a robust susceptible phenotype, while DIO-hM3Dq mice displayed resilient phenotypes following CNO drinking water administration (Fig. 3C). Moreover, none of the DIO-hM4Di + CNO mice went on to develop resilience (0/10, SI >1.0) while 89% (8/9, SI ≧1.0) of the DIO-hM3Dq were resilient (Fig 3C,F). Notably, the social defeat experience was not affected by the DREADDs manipulations (Supplemental Data Fig. 3A, B). The SP test — a test of hedonic behavior — revealed differences between mCherry susceptible (mCherry (S)) and resilient (mCherry (R)) mice that were mirrored in DIO-hM4Di and hM3Dq mice, respectively. mCherry (S) and hM4Di mice displayed a significant decrease in SP relative to mCherry (R) and hM3Dq-injected mice (Fig. 3D). There were no significant differences in locomotion (Fig. 3E). The chemogenetic manipulation also produced bidirectional effects on anxiety-like behavior in elevated plus maze (EPM) and open-field tests (Supplemental Data Fig. 4). Surprisingly, mice injected with hM3Dq-DREADDs went on to become resilient (Fig 3C) although activation of *Crf* neurons of the BNST have previously been shown to produce depressive- and anxiogenic-like responses.^27,34,39,46,49,57^ Interestingly, resilience was established if BNSTov^CRF^ chemogenetic activation occurred between 7 and 10 SDEs; excitatory hM3Dq-DREADDs activation during episodes 4-7 or 10-13 failed to display resiliency (Fig. 3G-I), suggesting stress history plays a critical role in the behavioral outcomes of CRF modulation. Modulating *Crf* neurons with inhibitory hM4Di- or excitatory hM3Dq-DREADDs between 8 and 10 SDEs led to enduring susceptible or resilient phenotypes, respectively, up to 6 weeks after CNO manipulation (Supplemental Data Fig. 6). These observations strongly support that the behavioral outcomes induced by the activation of BNSTov^CRF^ neurons are stress-history dependent.

**Figure 3.**
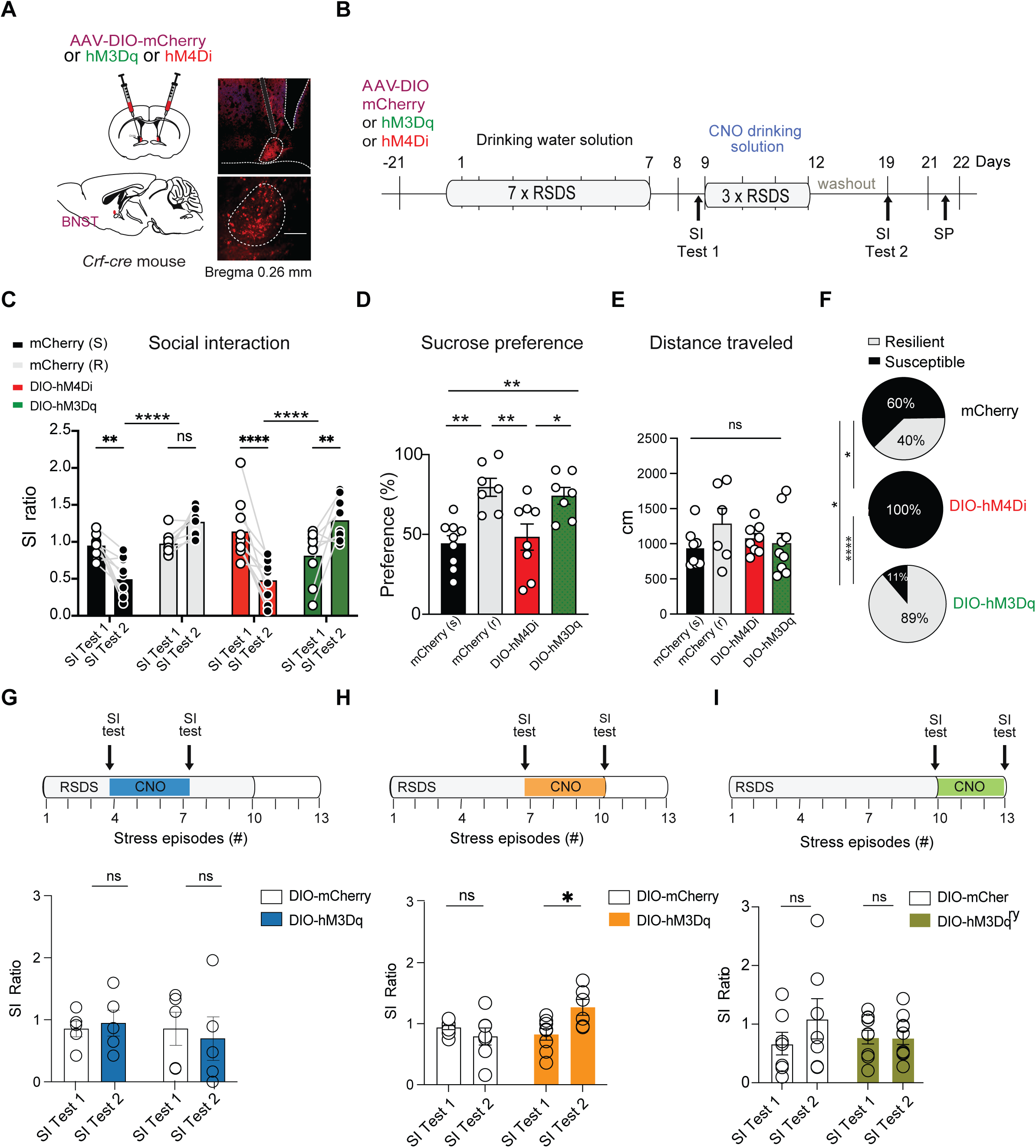
Chemogenetic modulation of BNSTov^CRF^ neurons bidirectional recapitulates behavioural tipping point. **A.** Viral targeting of BNSTov. **B.** Experimental design of chemogenetic manipulation of BNSTov neurons with CNO-drinking water construct. **C.** Social interaction test, two-way ANOVA F_(3,68)_=18.01, row ****P<0.0001, row factor F_(3, 68)_=9.965, P=0.2616 (time), row factor x time F_(1,68)_=1.281, ****P<0.0001, Sidak’s multiple comparisons test mCherry (s), **P=0.0017 (n=10 mice); mCherry (r), P=0.1322 (n=9 mice); hM4Di, ****P<0.0001 (n=10 mice); hM3Dq, **P=0.0018 (n=9 mice). **D.** Sucrose preference test. one-way ANOVA treatment F_(3,27)_=8.310, ***P<0.001, Tukey’s post-hoc testing susceptible vs hM4Di, P=9619; susceptible vs resilient, **P<0.01; susceptible vs hM3Dq, **P<0.01; hM4Di vs resilient, **P<0.01; hM4Di vs hM3Dq, *P<0.05; resilient vs hM3Dq P=0.9368 (7-9 mice per group). **E.** Distance traveled, one-way ANOVA F_(3,27)_=1.031, P=0.3944. **F.** Percentage of susceptible or resilient mice: DIO-hM4Di, 100% susceptible; DIO-hM3Dq, 89% resilient and 11% susceptible; mCherry, 40% resilient and 60% susceptible. Chi-square test=15.66, df=2, ***P<0.001 (n=21 mCherry, 10 hMDi, 9 hM3Dq); Fisher’s post-hoc test mCherry vs hM4Di, *P<0.05; mCherry vs hM3Dq, *P<0.05; hM4Di vs hM3Dq, ****P<0.0001. **G.** Repeated social defeat DREADDs manipulation (4-7 episodes of stress), 2-way ANOVA treatment F_(1,16)_=0.01749, P=0.8964; SI Test 1 vs SI Test 2, F_(1,16)_=0.2377, P=0.6325; interaction F_(1,16)_=0.07055, P=0.6218 (n=5-6 mice/group). **H.** RSDS DREADDs manipulation (7-10 episodes of stress), 2-way ANOVA treatment, F_(1,22)_=8.675, **P< 0.01; SI Test 1 vs SI Test 2, F_(1,22)_=2.549, P=0.1247; interaction F_(1,22)_=2.549, P=0.1247. Sidak’s post-hoc test SI Test 1 vs SI Test 2, control, P=0.5533; DIO-hM3Dq, *P<0.05 (n=6-7 mice/group). **I.** RSDS DREADDs manipulation (10-13 episodes of stress), 2-way ANOVA F_(1,30)_=1.192, treatment, P=0.2623. SI Test 1 vs Si Test 2, F_(1,30)_=0.3132, P=0.2836. Interaction F_(1,30)_=0.2623, P=0.5799. All data represent means ± SEM. *P<0.05, **P<0.01, ***P<0.001, ****P<0.0001, ns = not significant.

**Figure 4.**
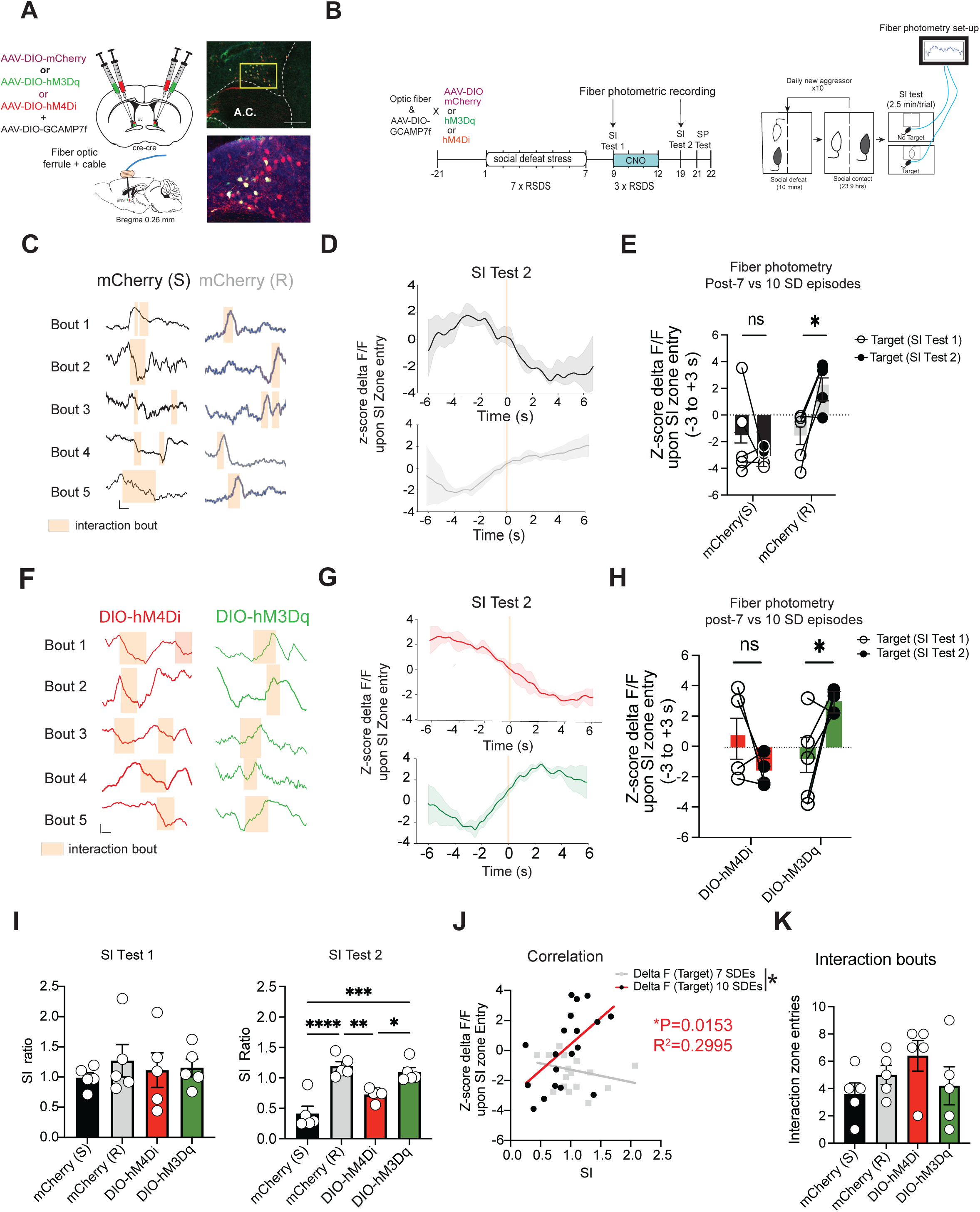
BNSTov^CRF^ calcium-dynamics encode stress effect on social interaction. **A.** Viral targeting of the BNSTov. **B.** Experimental design of multiplexed chemogenetics with drinking water-CNO delivery and fiber photometry. **C.F.** Representative calcium recordings of mCherry susceptible (S), mCherry resilient (R), hM3Dq, and hM4Di groups, respectively. Vertical scale bar is equal to z-score of 1, and horizontal scale bar is equal to 10 s. **D. G.** Representative averaged trace centered around interaction bout for the four groups stated above. **E.** Two-way RM ANOVA row factor F_(3,14)_=1.401; pre vs post-CNO F_(1,14)_=0.8179; subject F_(14,14)_=1.195; Row x SI Test 1/SI Test 2, F_(3, 14)_=9.343, **P<0.01. Sidak’s post-hoc test, susceptible vs resilient, P=0.137 vs *P=0.05, respectively (n=4 mice/group). **H.** hM4Di vs hM3Dq, P=0.3069 vs *P<0.05, respectively (n=4-5 mice/grp). **I.** SI Test 1 social interaction test. One-way ANOVA F_(3,16)_=0.291, P=0.8308 (n=5 mice/group). SI Test 2 social interaction test, One-Way ANOVA F_(3,16)_=17.83, ****P<0.0001; Tukey’s post-hoc test, susceptible vs resilient, ****P<0.0001; susceptible vs hM3Dq, ***P<0.001; susceptible vs hM4Di, P=0.0762; resilient vs hM3Dq, P=0.8506; resilient vs HM4Di, **P<0.01; hM3Dq vs hM4Di, *P<0.05 (n=5 mice/group). **J.** Correlation of SI ratio and z-score delta F/F upon social entry, simple linear regression pre-CNO F_(1,15)_=0.7674, P=0.3948; SI test 2 F_(1,17)_=7.268, *P<0.05. Intersection of lines, F_(1,32)_=6.896, *P<0.05 (n=18 mice). **K.** Number of social interaction bouts, one-way ANOVA F_(3,16)_=1.346, P=0.2949 (n=5 mice/group). All data represent means ± SEM. *P<0.05, **P<0.01, ***P<0.001, ****P<0.0001, ns = not significant.

### Calcium dynamics underlying stress adaptation

While we showed that we could bidirectionally drive the establishment of resilience, this manipulation offers limited insight into the *in vivo* neural dynamics that reflect this divergence point. To draw a link between neural changes and behavior *in vivo*, we combined fiber photometry (gCAMP7f) with excitatory or inhibitory DREADDs (Cre-dependent -hM4Di, -hM3Dq DREADDs or -mCherry viral vectors) to mimic the neuroadaptive changes in BNSTov^CRF^ and bidirectionally drive the development of resilience/susceptibility (Fig 4A, B). Clozapine-N-oxide (CNO) was administered via drinking water over the last 3 episodes of RSDS.

Given that mice with shared stress-history of 7 SDEs appeared to have unimodal distribution on a variety of measures, we hypothesized that calcium-encoded neural activity may diverge in susceptible/resilient mice concomitant with the display of their respective behaviors on SI test. When comparing changes in calcium-encoded neural dynamics between SI Test 1 and 2, mCherry (S) mice showed no difference, whereas mCherry (R) mice developed an increase concomitant with the display of resilience (Fig. 4C-E), not observed in mice enduring only 7 SDEs. These findings suggest that the persistence of neural activity across SDEs #8-10 is associated with resiliency.

During SI Test 1, similar to control mice, DREADDs injected mice experienced a decreased in calcium-related neural activity upon initiating social interaction with a novel conspecific (Supplemental Data Fig 7A), whose neural pattern was observed in both no target and target trials (Supplemental Data Fig 7B). In hM4Di-injected mice, there were no significant differences in neural activity during SI Test 1 and 2 trials, similar to control mCherry (S) mice (Fig. 4F-H; Supplemental Data Fig. 7A-C). As was observed in the resilient (mCherry (R)) mice, the hM3Dq-injected group showed an increase in calcium-encoded neural activity (Fig 4F-H). Mice injected with cre-dependent hM4Di showed a significant decrease in calcium-related neural activity. In contrast, social interaction initiation led to increased activity in DIO-hM3Dq injected mice (Fig. 4F-H). There were no significant differences in SI of mice subjected to 7 SDEs, yet differences emerge following 3 additional SDEs with a subset of mice becoming susceptible and resilient (mCherry(R)/(S)) or activating/inhibiting DREADD promoting resiliency and susceptibility in the direction predicted (Fig. 4I). We observed a strong correlation between SI ratio and calcium activity only after 10 SDEs (Fig. 4J). There were no significant differences observed in calcium-based neuronal activity upon social interaction with a novel conspecific in mice subjected to 7 SDEs (Supplemental Data Fig. 7A,B), or in the absence of social interaction (Supplemental Data Fig. 7C). In contrast, after 10 SDEs, mCherry (R) and DIO-hM3Dq injected mice displayed significantly greater neuronal activation upon social contact relative to mCherry (S) and DIO-hM4Di injected mice (Supplemental Data Fig. 7D). Despite the difference in time spent interacting with a novel conspecific, the number of interaction zone entries were not significantly different, nor differences in distance traveled (Fig. 4K, Supplemental Data Fig. 7E). These data suggest that BNSTov^CRF^ neuronal dynamics are differentially altered by stress modulation in accordance with phenotypical display resiliency/susceptibility.

### *Crfr1* expression in BNSTov^CRF^ neurons mirrors the behavioral emergence of resilience

We observed a stress-induced enhancement in firing rates of BNSTov^CRF^ neurons in resilient mice. To explore the effect of this BNSTov^CRF^ stress modulation on CRF receptor transmission, we used RNAScope *in situ* hybridization to quantify *Crfr1* and *Crfr2* in accordance with stress history. Mice were subjected to either 7- or 10-daily episodes of social defeat, and the BNST and *Crfr1* and *Crfr2* genes were assessed (Fig. 5A-C) due to their reported role in mediating stress responses^30,58–60^. The percentage of CRFR1-expressing neurons among CRF-expressing neurons was higher in mice subjected to 7 SDEs than in susceptible mice but not significantly different than resilient mice (Fig. 5D). In contrast, there were no significant differences in BNSTov neurons co-expressing *Crfr2 and Crf mRNA* across groups of mice (Fig. 5E). The overlap between *Crfr1* and *Crf* in the BNST was significantly greater in the oval nucleus than in anterolateral, anteromedial, and ventral subregions of the anterior dorsal BNST (Fig. 5C,F).

**Figure 5.**
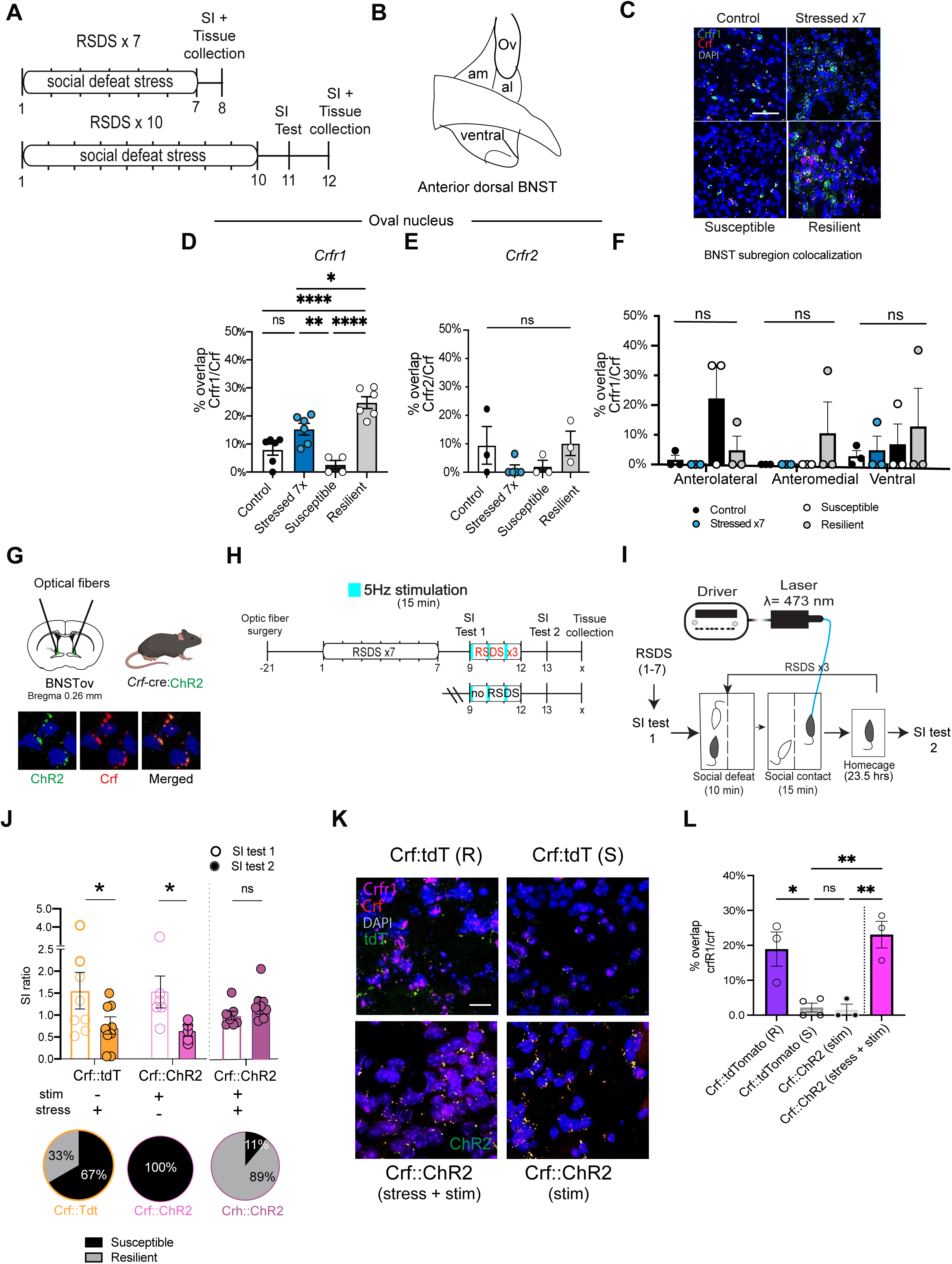
BNSTov *Crfr1* is associated with the emergence of resiliency. **A.** Experimental timeline of RNAScope *in situ* hybridization of mice that underwent social defeat stress. **B**. Representative images of control, stressed x7, susceptible, and resilient. 20x magnification, scale bar 0.64 mm. **C.** Schematic of anterior dorsal BNST. **D.** *Crfr1* mRNA colocalization, one-way ANOVA F_(3,18)_=20.91, ****P<0.0001; Tukey’s posthoc test, control vs stressed x7, P=0.0701; control vs susceptible, P=0.3336; control vs resilient, ****P<0.0001; stressed x7 vs susceptible, **P<0.01; stressed x7 vs resilient, *P<0.05; susceptible vs resilient, ****P<0.0001 (n=4-6 BNST brain samples). **E.** *Crfr2* mRNA colocalization, one-way ANOVA F_(3,10)_=1.790, P=0.2166 (n=3 BNST brain samples). **F.** RNAscope BNST subregion comparison, two-way ANOVA interaction F_(6,24)_=1.057, P=0.4147; row factor F_(2,24)_=0.6092, P=0.5520; column factor F_(3,24)_=1.549, P=0.2275 (n=3 BNST brain samples). **G.** Viral injection site. **H.** Experimental timeline. **I.** Optogenetic and social behavioral setup. **J.** Optogenetics social interaction, two-way ANOVA interaction F_(1,28)_=5.059, *P<0.05; row factor F_(1,28)_=1.061e-005, P=0.9974; column factor F_(1,28)_=1.419, P=0.2436. Sidak’s post-hoc test, *Crf*::tdT, *P<0.05; *Crf*::ChR2, P=0.7233 (n=7-9 mice/group). Optogenetic manipulation unpaired t-test, *Crf*::tdT, t=2.556, df=40, *P<0.05 (n=8-9 mice); *Crf*::ChR2, t=0.7758, df=40, P=0.442457 (n=8-9 mice); *Crf*::ChR2, t=2.369, df=40, *P<0.05 (n=6 mice). Holm-Sidak method for multiple comparisons. **K.** Representative image, scale bar 0.64mm. **I.** Optogenetics-RNAScope experiment. One-Way ANOVA F_(3,9)_=13.53, **P<0.01. Tukey’s post-hoc test, *Crf*::ChR2 (stress + stim) vs *Crf*::tdT (resilient), P=0.7872; *Crf*::ChR2 (stress + stim) vs. *Crf*::tdTomato (s), **P<0.01; *Crf*::ChR2 (stress + stim) vs. *Crf*::chR2 (stim), **P<0.01; *Crf:*:tdTomato (r) vs. *Crf:*:tdTomato (s), *P<0.05; *Crf*::tdTomato (r) vs. *Crf:*:ChR2 (stim), *P<0.05; *Crf*::tdTomato (s) vs. *Crf*::ChR2 (stim), P=0.9988. All data represent means ± SEM. *P<0.05, **P<0.01, ***P<0.001, ****P<0.0001, ns = not significant.

To explore the role of firing rate changes on gene expression and the development of resiliency, we optogenetically stimulated BNSTov^CRF^ neurons using transgenic *Crf-Cre*::ai32 mice, which express channelrhodopsin-2 (ChR2) in *Crf*-containing neurons ^61^ (Fig. 5G). For control experiments, transgenic mice of a similar background were used, except instead of ai32, mice expressed the tdTomato fluorophore in CRF neurons (*Crf-tdTomato*). 5 Hz stimulation frequency was used in order to mirror the average firing rate observed in the spontaneous firing rate of resilient mice (Fig. 2D). Mice received 15-minute 5 Hz photo-stimulation of BNSTov^CRF^ neurons following physical stress on SDEs #8-10. Mice were placed in the adjacent compartment separating the aggressor, preventing physical contact. The compartment divides the aggressor cage in half by a clear plexiglass that allows for continuous sensory cues (Fig. 5H,I). Photo-stimulation of *Crf*::ChR2 mice led to a significantly higher SI ratio and greater percentage resilient than *Crf::tdTomato* mice (89% vs. 33%) when stimulated during SDEs #8-10 (Fig. 5J). Surprisingly, *Crf::ChR2* mice that received photo-stimulation but were not subjected to SDEs #8-10 showed a significant decrease in SI ratio and 100% of the mice, becoming susceptible (0/6, SI ≧ 1.0) (Fig. 5J). Photo-stimulation paired with SDEs #8-10 increased the percentage of cells co-expressing *Crfr1* mRNA in CRF+ neurons relative to mice that experienced photo-stimulation in the absence of additional stress (Fig. 5K, L). Additionally, we obtained slice preparation from the optogenetically-induced resilient mice, and performed cell-attached single unit recordings from BNSTov ChR2-expressing CRF neurons (Supplemental Data Fig. 8A). Our recording data showed that optogenetic stimulation reliably induced 2.5 Hz, 5 Hz and 10 Hz spike responses as expected (Supplemental Data Fig. 8B), and interestingly triggered burst firing after the 5 Hz optical stimulation (Supplemental Data Fig. 8C,D). Importantly, Bath applied CRFR1-selective antagonist NBI 27914 significantly decreased the firing rate of tested BNSTov ChR2-expressing CRF neurons (Supplemental Data Fig. 8E,F).

In summary, these data show that maintenance of resiliency requires social defeat stress BNSTov^CRF^ activation and is correlated with the upregulation of *Crfr1* expression in a stress-history dependent manner.

## Discussion

The RSDS paradigm was modified to observe the effects of cumulative stress on neuroplasticity in regions critical for mood regulation. Indeed, our work has uncovered a discrete window of neuronal and behavioral plasticity between 7 and 10 SDEs, during which susceptible and resilient phenotypes are established. By capturing behavioral, electrophysiological, and *in vivo* fiber photometric measures during the intra-social defeat stress period, we uncover the mechanisms underlying the establishment of resilience.

The oval nucleus of the BNST has been investigated in stress responses, we hypothesized that the region would be instrumental in processing contexts associated with social stress. Indeed, we observed that individual differences of stress effects on social behavior are encoded by BNSTov^CRF^ neurons. Prior studies have implicated the overactivation of CRF neurons in the BNST as being pro-depressive and anxiogenic^31,39,46,47,49^, therefore, we were surprised to observe that BNSTov^CRF^ neurons exhibited increased activity in resilient mice. When these same cells were active, they yielded resilient mice. Indeed, we observed that Cre-dependent hM3Dq action produced resilient mice only after a certain “dose” of daily stressors (7 SDEs), suggesting that the BNSTov may be tightly modulated based on stress history. Our findings support a view that resilience is a stress history-dependent state, which does not exist before stress. This is highly consistent with previous demonstrations that resilience is a status achieved by active regulation of genes and ion channel functions in this group more than in susceptible animals^20–21,62–67^.

CRF neurons have been observed to influence the salience of stressful contexts according to stress exposure^24,27,60,64,68^. One feasible mechanism by which this occurs is through CRF receptor dynamics. We hypothesized that the development of resilience occurs via CRF-CRFR interactions in the BNST. Surprisingly, we observed *Crfr1* mRNA expression in CRF (and not neighboring CRF^-^) neurons, suggesting that stress works as a pro-resilient agent based on stress-history. This is in contrary to what has been observed, namely that CRFR1 has been found largely on non-CRF neurons in the BNST.^68,69^

CRFR1 is selectively activated in the context of ongoing stress, serving as a coincidence-detector ^70^. Chronic stress has been shown to shift the connectivity of local CRF^+^ neurons from CRF^+^-CRF^-^ to a larger percentage of CRF^+^-CRF^+^ cells^73^. Studies using prolonged overactivation (over weeks to months) of CRF activity have yielded antidepressant and anxiolytic results^29,34,35^. Here, we identify a more discrete timeline in the span of days to capture the transition of when BNSTov^CRF^ activation becomes pro-resilient. By optogenetically activating BNSTov^CRF^ neurons, we observed an increase in CRFR1 expression. Though correlative, this exquisite regulation of CRFR1 according to stress-history and at times demonstrating an opposing effect on depressive-like behavior, may underlie why clinical trials of CRFR1 antagonists for MDD have been met with variable success^59,74–76^. Although beyond the scope of the current study, future experiments using siRNA or CRISPR knockdown approaches should be performed to determine whether upregulation of CRFR1 is a critical facet of neuroadaptation leading to resiliency.

Stress-sensitive regions such as the BNST have been found to be of critical importance in stress coping and reactivity^77–79^. Stress resilience has long been considered a response separate from or in the absence of stimuli that gives rise to stress susceptibility, mediated by parallel circuits or cell-types in a particular brain region^14,15,80^. Here, we observe that activity dynamics of CRF neurons can shape and influence resiliency to stress, potentially through (auto)regulation of *Crfr1* mRNA. Other substrates may be used to classify these neurons either on the basis of dual-neuropeptide or circuit-specific identities and should be the object of future exploration.

Previous work has shown that resiliency is influenced by dopaminergic VTA neurons in the nucleus accumbens (NAc)^29^, in part, through the actions of brain-derived-neurotrophic-factor (BDNF)^21,81,82^. CRF peptide has been important for BDNF release in the NAc as a stress-coincidence sensor^20^, yet the sources of CRF important for altering stress effect on social and hedonic behavior has not been extensively characterized. While long-range GABAergic BNST neurons projecting to the VTA have been shown to influence reward and anxiety-like behavior^48,83–86^, it is unclear to what degree these cells comprise the oval nuclear BNST population. The BNST also sends projections to the dorsal raphe, lateral and paraventricular hypothalamus, and ventrolateral periaqueductal gray^27,31–33,42,77,87–89^, among others, whose modulation have been linked to stress on affect and social motivation. In this way, the BNST acts as a node for integrating information regarding stress history and determining socio-affective outcomes according, possibly due to CRFR1 receptor dynamics occurring on CRF neurons of the oval nucleus, thereby shaping the long-lasting outcome of resiliency.

Our study highlights a previously unknown mechanism by which the BNST encodes cumulative social stress and effectuates susceptible or resilient outcomes. Importantly, there are currently no Food and Drug Administration-approved drugs aimed at preventing a depressive episode from occurring. By targeting mechanisms involved in establishing resiliency, the possibility may exist to therapeutically leverage windows of plasticity to effectuate resilience and evade the development of MDD.

### Statistical analysis

Animals were randomly assigned to control and experimental groups and all experimenters were blinded. Mice were excluded if viral infection was off-target. No data was excluded for other reasons. Student’s two-tailed t-tests were used for comparisons of two experimental groups. For parametric data sets, comparisons among three or more groups were performed using one or two-way ANOVA tests followed by Tukey’s or Bonferroni posthoc tests. For all tests, p<0.05 was determined to be significant. Statistical analyses were performed using Graph Pad Prism 9 .3.1 software (La Jolla, CA, USA). For data not normally distributed, non-parametric analyses were performed.

## Data & Code availability

All source data for this manuscript and MATLAB code used to analyze photometry data is available upon request.

## Acknowledgements

We would like to thank S.M. for generous sharing for input on the manuscript. We would like to thank S.R. and L.L. for lending viral constructs for behavioral validation. We would like to thank the animal care staff at ISMMS for the animal care staff and technical support. This research was funded by the National Institutes of Health grants R01MH072908 to DGR and LJY, R01MH1206387 to MHH and SJR, R21MH112081 to MHH and SJR, F31MH114624 to SEH, and P51 OD011132 to YNPRC, by the National Key R&D Program of 1034 China No. 2021ZD0202900 to MHH.

