## Supplementary Methods for "Stress History Modulates CRF Neurons to Establish Resilience"

**Mice**

Upon receipt from Jackson Laboratories, mice were acclimated to the housing facility for 1-2 weeks prior to the start of experiments. All mice were group-housed and maintained on a 12/12 light-dark cycle with ad libitum access to food and water. Following social defeat stress, mice were singly housed and maintained on a 12/12 light-dark cycle with ad libitum access to food and water.

**Stereotaxic virus and optic fiber implantation**

Under ketamine (80 mg/kg)/xylazine (10 mg/kg) anesthesia, mice were placed in a stereotaxic frame (Kopf Instruments) and the BNSTov was targeted (coordinates: anterior/posterior +0.20, media/lateral: +/- 2.15, dorsal/ventral, –4.0 mm; 15-degree angle). Hair was shaved around the crown of the head, alcohol, and betadine were applied to the scalp. Ophthalmic ointment was applied to the eyes to prevent dryness, a midline incision was made down the scalp, and a craniotomy was made using a dental drill. A 10 ul Nanofil Hamilton syringe (WPI, Sarasota, Fl) with a 34-gauge beveled metal needle was used to infuse 0.5 ul virus at a rate of 85 nl/minute. Following infusion, the needle was kept at the injection site for 10 minutes and slowly withdrawn. For the optogenetic experiments, chronically implantable optic fibers constructed with 1.25 mm diameter, 200 um core, 0.39 numeral aperture (NA) were used (RWD). For the fiber photometry experiments, chronically implantable optic fibers constructed with 1.25 mm diameter, 400 um core 0.48 numerical aperture (NA) optic fiber and unilaterally implanted into the BNSTov (coordinates: anterior/posterior +0.20, media/lateral: +/- 2.15, dorsal/ventral, –3.9 mm; 15º angle) (thor labs). Fiber optical ferrules were cemented to the skull using dental acrylic (Parkell C&B Metabond). All optical stimulation and fiber photometric recording experiments were conducted a minimum of 3-4 weeks post-implantation.

**Optogenetic manipulation of BNSTov^CRF^ neurons**

Optical fibers were implanted on Ai32 (transgenic mouse line expressing light-gated cation channel channelrhodopsin-2 (ChR2). Optical fiber Optogenetic stimulation was conducted via the use of a diode-pumped solid-state (DPSS) 473-nm blue laser (Crystal Laser, BCL-473-050-M), using a patch cord with an FC/PC adaptor (Doric Lenses, MFP_200/240/900-0.22_4m_FC-MF2.5). A functional generator (Agilent Technologies; 33220A) was used to generate a 5 Hz frequency, pulse width of 10 ms for 15 minutes[^97,98^](https://sciwheel.com/work/citation?ids=1955437,883116&pre=&pre=&suf=&suf=&sa=0,0&dbf=0&dbf=0), and power density between 7-9 mW mm^-2.^ Experimenters were blinded to the stimulation group.

**Fiber photometry calcium imaging**

Optical recordings of GCaMP7f fluorescence were acquired using two LEDs at 490 and 405 (Thor Labs), reflected off dichroic mirrors (Semrock, FF495) and coupled into a 400 micro 0.48 NA optical fiber (Thorlabs BFH48-600) using a 40 x 0.48 NA microscope objective and fiber launch with the pat chord linked to an implanted 400 um optical fiber with zirconia sheath. Signals in both 470 and 405 nm channels are monitored throughout the recordings, whereas the 405 nm is used as an isosbestic control for ambient fluorescents and motion artifacts caused by movement or torque about the fiber optic implant. Wavelengths were modulated at frequencies of 210-220 and 330 Hz, respectively, with power output maintained at 20 mA and a DC offset of 3 mA for both light sources. All signalers were acquired at 1 kHz and lowpass filtered at 3 Hz. Prior to social interaction testing, mice were handled for and connected via a patch cable and placed in the home cage for 5-7 minutes for basal BNSTov recording and habituation prior to the start of the no-target trial.

**Fiber photometry analysis**

A custom-written MATLAB code (modified from previously published work)[^99^](https://sciwheel.com/work/citation?ids=2951165&pre=&suf=&sa=0&dbf=0) was used to analyze the GCaMP7f signal. Firstly, the bulk fluorescent signal from both the 470 nm and 405 nm channels were normalized. A linear regression (slope of the 405 nm fitted against the 470 nm signal) was applied over the data to correct for bleaching of signal of the duration at each recording. The Initial 3 seconds for the signals were discarded because of the photoreceiver/LED rise time artifact. Detection of GCaMP7f signal is calculated as a change in the 470 nm/fitted 405 nm signal over the fitted 405 signal (ΔF/F). For the calculation of intensity values around events of interest (PC2), a 6-second window was used. We used 3 seconds before and after the PC2 event onset and computed z-scores for all such event windows. These z-scored intensity values were then averaged within and between animals in particular groups, and 95% confidence intervals for the averaged intensities were computed.

**Cell-attached electrophysiology**

Acute coronal brain slices of the BNSTov were prepared according to previously published protocols[^65,66,90,94^](https://sciwheel.com/work/citation?ids=6005676,123261,663599,81187&pre=&pre=&pre=&pre=&suf=&suf=&suf=&suf=&sa=0,0,0,0&dbf=0&dbf=0&dbf=0&dbf=0). All recordings were carried out blind to stress history, behavioral phenotype, or drug treatment. Male 8-12 weeks old mice were perfused with cold artificial cerebrospinal fluid (aCSF) containing (in mM): 128 NaCl, 3 KCl, 1.25 NaH_2_PO_4_, 10 D-glucose, 24 NaHCO_3_, 2 CaCl_2_, and 2 MgCl_2_ (oxygenated with 95% O_2_ and 5% CO_2_, pH 7.35, 295-305 mOsm). Acute brain slices containing BNSTov were cut using a microslicer (DTK-1000, Ted Bella) in sucrose-ACSF, derived by replacing NaCl with 254 mM sucrose, and saturated by 95% O_2_ and 5% CO_2_ (2.5 ml/min) and 35ºC. Glass recording pipettes (2-4 MΩ) were filled with an internal solution containing (mM): 115 potassium gluconate, 20 KCl, 1.5 MgCl_2_, 10 phosphocreatine, 10 HEPES, 2 magnesium ATP and 0.5 GTP (pH 7.2, 285 mOSm). BNSTov CRF neurons were identified by location, and infrared differential interference contrast microscopy and recordings were made from CRF-positive neurons as indicated by the presence of tdTomato (in *Crf*::tdTomato mice) or eYFP in the case of (*Crf*::ChR2) mice. Spontaneous firing rates were recorded in cell-attach mode, and data acquisition and online analysis of firing rates were collected using a Digidata 1440 digitizer and pClamp 10.2 (Axon Instruments). The timing of recordings was made consistent throughout treatment groups. A burst was defined as containing at least three spikes, with interspike intervals <15 ms[^95^](https://sciwheel.com/work/citation?ids=9165366&pre=&suf=&sa=0&dbf=0). Neuronal firing rates were considered bursting or non-bursting if they had undergone a statistically significant change with a P<0.05 on the rank-sum test.

**Repeated Social Defeat Stress (Continued):**

During the period which marks the physical stress, C57BL/6J mice are placed on the ipsilateral side of the cage as the CD1 aggressor for 10 minutes. Following this, the intruder mouse is placed in the contralateral side of the perforated Plexiglas divider for the remainder of the 24-hour period, marking the sensory-stress period. Every 24 hours, the intruder mouse is paired with a new aggressor for 10 episodes. Control mice were housed two mice per cage divided by a perforated Plexiglas divider and rotated and handled daily like the socially defeated mice.

**Social interaction test**

Social interaction testing was performed as described[^6,20,21,65,66,81,82,93^](https://sciwheel.com/work/citation?ids=72311,7722735,1230587,73234,3174591,6005676,123261,1135433&pre=&pre=&pre=&pre=&pre=&pre=&pre=&pre=&suf=&suf=&suf=&suf=&suf=&suf=&suf=&suf=&sa=0,0,0,0,0,0,0,0&dbf=0&dbf=0&dbf=0&dbf=0&dbf=0&dbf=0&dbf=0&dbf=0). Briefly, a novel conspecific of CD1 strain is placed in an interaction zone of a standard open-field arena, and the time the intruder spends in the interaction zone is measured.  The mice spend a total of 5 minutes in the open arena (2.5 min with and without the novel CD1). Ethovision (Noldus Information Technology) video-tracking software is used to track interaction time. All social interaction testing takes place 24 hours after the last defeat. Social interaction (SI) is measured by time spent in the interaction zone during first (CD1 absent) over second (CD1 present) trials. Mice are categorized according to SI ratios; an SI ratio ≧1 defines resilient, whereas an SI ratio <1 is susceptible as described previously[^5,17,19,21^](https://sciwheel.com/work/citation?ids=72311,6526600,12243797,12206169&pre=&pre=&pre=&pre=&suf=&suf=&suf=&suf=&sa=0,0,0,0&dbf=0&dbf=0&dbf=0&dbf=0).

**Sucrose preference test**

For sucrose preference testing, a solution of 1% sucrose or diluent alone (drinking water) is filled in 50 ml tubes with ball-pointed sipper nozzles (Ancare). Animals are acclimatized to two-bottle choice conditions prior to testing. The bottles are weighed, and positions interchanged daily. Sucrose preference is calculated as a percentage [100 x volume of sucrose consumed (in bottle A)/total volume consumed (bottles A and B)] and averaged over 2 days of testing.

**Qualitative defeat assessment**

Social defeat encounters were recorded, and BORIS (Behavioral Observation Research Interactive Software) was used to code behaviors. Five behaviors (cage exploration, aggressor grooming, flight, motionless, rearing/defensive posturing) were scored per published reports on typical social defeat behaviors[^9,18,37,96^](https://sciwheel.com/work/citation?ids=4809818,4872691,12243933,12378472&pre=&pre=&pre=&pre=&suf=&suf=&suf=&suf=&sa=0,0,0,0&dbf=0&dbf=0&dbf=0&dbf=0). The total time engaging in each behavior was tallied to a blinded observer, and scores were aggregated.

**CNO-drinking water construct**

Clozapine-N-Oxide (CNO) was obtained from (Hello Bio, catalog no HB6149). The dry chemical was dissolved in drinking water obtained from the vivarium and diluted such that each mouse received 5 mg/kg/day based on previous studies. CNO was made fresh each day for the three days it was administered. CNO solutions were protected from light throughout the experimental procedure. On average, mice consumed ~4-5 mL of water per day. Water bottles and mice were weighed daily. CNO water bottles were exchanged for normal drinking water after the social interaction test 1 and 8-12 hours before social defeat stress to ensure neuronal modulation during stress exposure.

**Elevated plus-maze**

Mice began testing by being placed in the center of the maze (Model ENV-560A, Med Associated, Fairfax, VT, USA), where movement was tracked using Ethovision behavioral tracking software (Noldus). The test lasted 5 minutes.

**Open field test**

Mice were placed in an arena (42 cm (w) x 42 cm (d) x 42 cm (h); Nationwide Plastics, custom order). Mice were tracked using behavioral tracking software (Ethovision, Noldus), and time spent in the designated center and surround zone, as well as locomotor activity, were measured.

**RNAScope *in-situ* hybridization**

To prepare frozen sections, mice were placed in an air-tight chamber with 2.5 ml of isoflurane-infused cotton balls. After 45-60 seconds (after respiratory depression and loss of consciousness as evidenced by the negative tail and paw pinch), brains were acutely harvested and placed in a -80°C storage chamber until RNAScope *in-situ* hybridization protocol commenced. 16-μm coronal sections were collected on a cryostat (Leica Biosystems) at -20°C and mounted directly onto ColorFrost Plus microscope slides (Fischer Scientific). Slides were stored at -80°C until *in situ* hybridization (ISH) processing. ISH was performed using the RNAscope multiplex fluorescent kit (Advanced Cell Diagnostics, ACD) according to the manufacturer's instructions. The tissue was fixed in 4% paraformaldehyde (PFA) in PBS chilled to 4°C for 15 minutes, washed twice briefly in 1x PBS, then dehydrated in 50% ethanol, 70% ethanol, and twice in 100% ethanol for 5 minutes each at room temperature (RT). A hydrophobic barrier was traced around sections of interest using an ImmEdge pen (Fischer Scientific), and once the barrier dried, sections were incubated in Protease IV reagent for 30 minutes at RT, then rinsed twice in 1x PBS for 5 minutes. Probes for CRH and either CRHR1 or CRHR2 (ACD) were warmed, mixed, and placed onto sections for hybridization at 40°C for 2 hours in a HybEZ II oven (ACD), which was used for all subsequent incubations. Sections were washed twice in 1x wash buffer (ACD) for 2 minutes each, then incubated with a series of amplification reagents at 40°C, Amp 1-FL for 30 minutes, Amp 2-FL for 15 minutes, Amp 3-FL for 30 minutes, and Amp 4-FL Alt A or C for 15 minutes, washing twice in 1x wash buffer for 2 minutes each between steps. DAPI was applied to the sections for 30 s, then immediately coverslipped with ProLong Gold mounting medium (Invitrogen). Images were collected on a Zeiss LSM 780 confocal microscope at 20x, 40x, or 63x magnification. mRNA puncta were quantified manually using FIJI 1.0 software (Image J), all experimental conditions were blinded to the imager, and all cell analysis and quantification.

**Immunohistochemistry**

Mice were quickly anesthetized with urethane and perfused with cold 1x-PBS (OmniPure, Bio-Rad) followed by cold 4% PFA (Fisher Hamilton Scientific). Brains were placed in PFA at 4ºC for at least 24 hours, followed by being replaced with a 15% sucrose/PBS solution at 4ºC for 24 hours and brain sinking. The next day, brains were placed in a 30% sucrose/PBS solution at 4ºC for 24 hours. Once the brains sunk, it was washed in 1xPBS three times and mounted on a freezing microtome for sectioning at 35 micrometers.

Sections were washed three times for 10 minutes in 1x PBS, then blocked in 5% bovine serum albumin (Sigma-Aldrich) in 1x PBS with 0.3% Triton-X (Sigma-Aldrich) for 1 hour. Tissues were incubated with primary antibodies for rabbit anti-mCherry ((1:1000), Invitrogen), rabbit anti-c-Fos antibody (1:250, Milipore C-10, sc-271243) and goat anti-GFP (1:500, abcams) overnight at 4ºC. Sections were washed three times with 1xPBS and blocked with 0.3% Triton-X for one hour. Tissue was blocked with secondary antibodies goat anti-rabbit/568 (1:1000, Abcams) and donkey anti-goat Alexa-488 (1:500, Abcams) at room temperature for 1 hour. Sections were washed three times with 1x PBS for 10 minutes and mounted on slides with ProLong Gold antifade reagent with DAPI (invitrogen, P36931). Z-stacked images were acquired with a Zeiss LSM780 multi photon confocal system and z-stacks were collated using ImageJ (Fiji). Cell counters were obtained using the Cell Counter feature on ImageJ.
