## Supplementary Figures for "Stress History Modulates CRF Neurons to Establish Resilience"

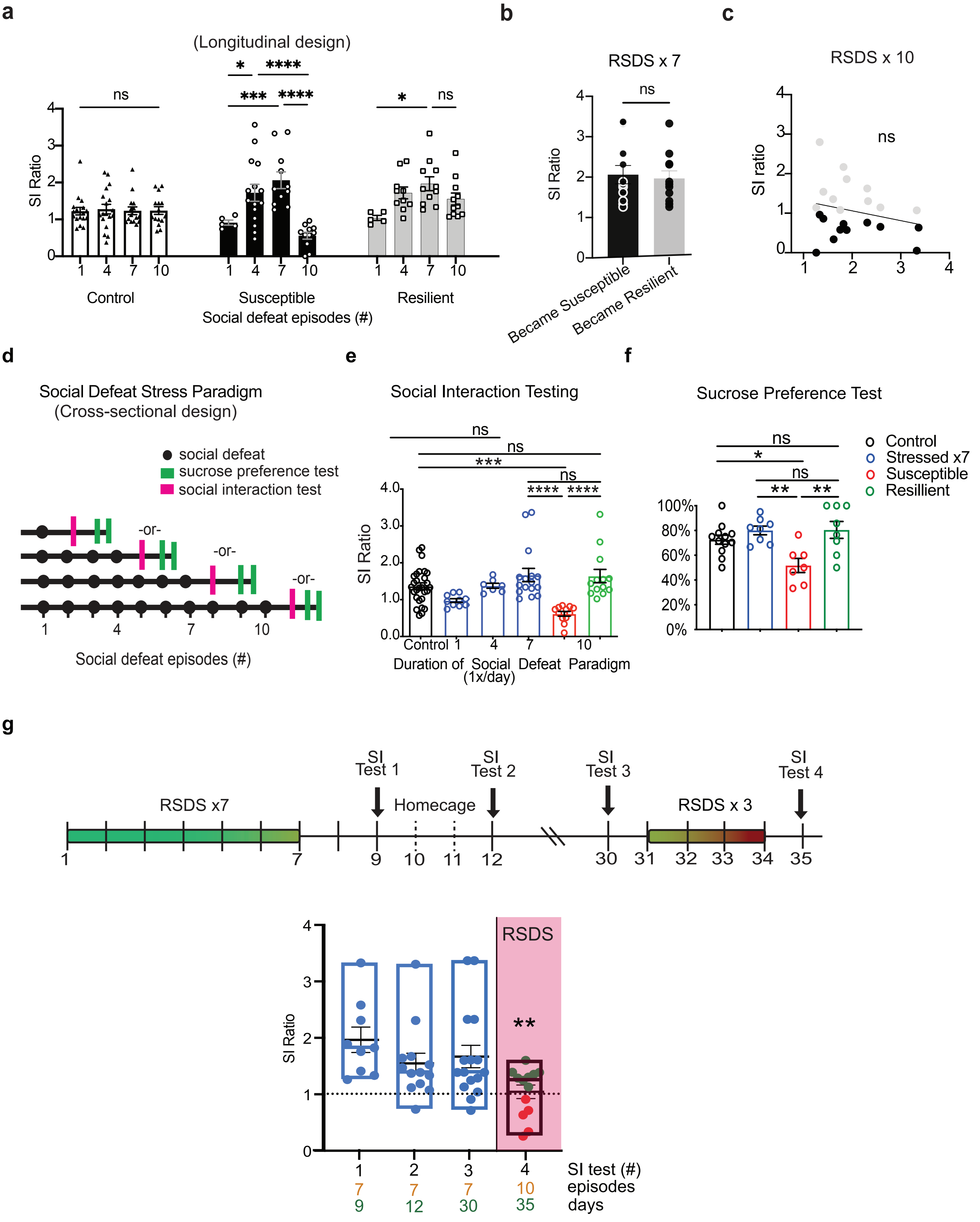


Social defeat episodes, not days, promulgate the divergence of susceptible/resilient phenotypes.

Extended Data Figure 1

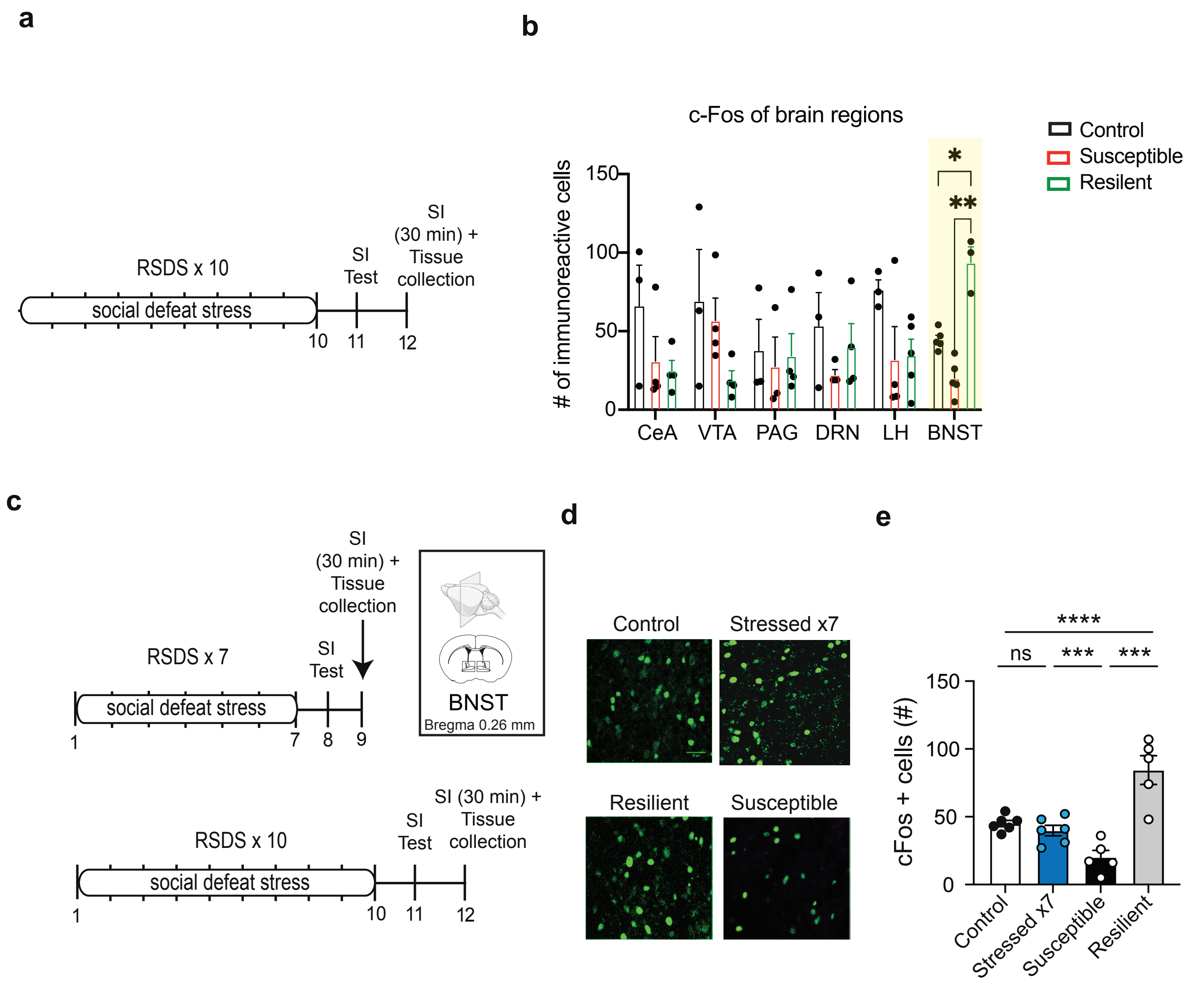


cFos immunohistochemistry of stress-sensitive regions reveals distinct activity patterns in the BNST during social interaction

Extended Data Figure 2

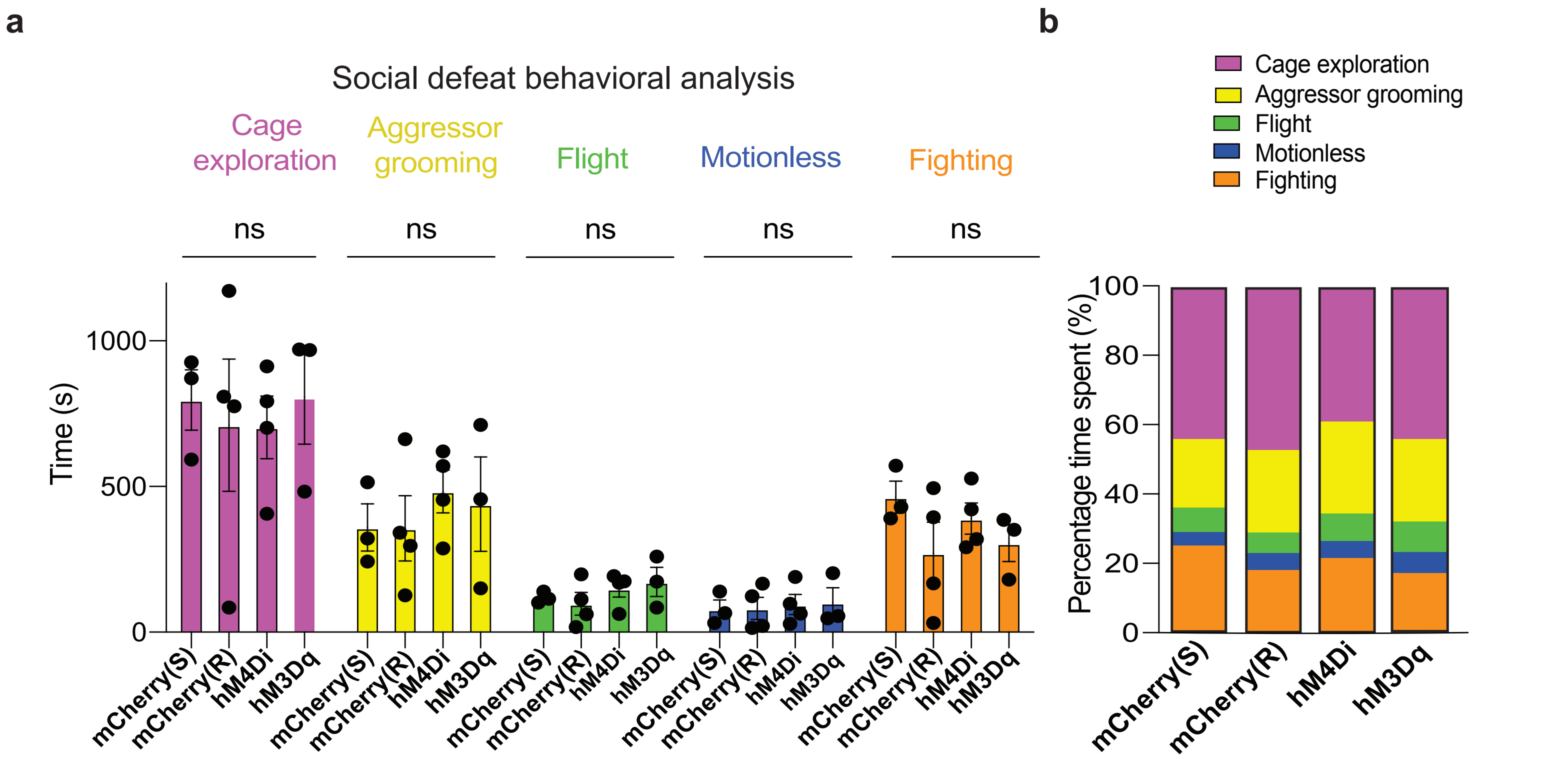


Chemogenetic manipulation of BNSTovCRF neurons does not effect social defeat dynamics between aggressor CD-1 and C57/BL6J subordinate mice.

Extended Data Figure 3

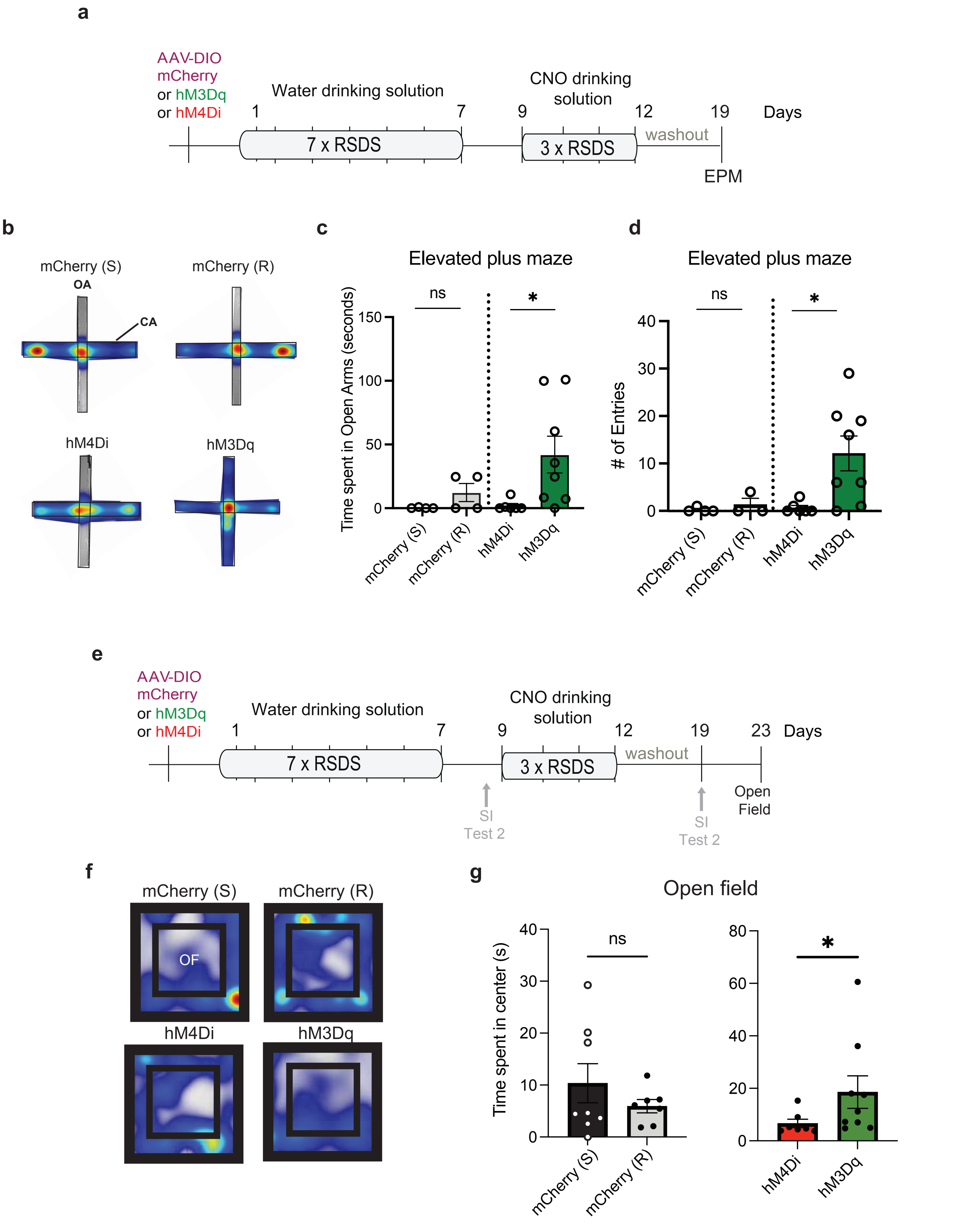


Chemogenetic manipulation of BNSTov CRF neurons bidirectionally modulates anxiety states

Extended Data Figure 4

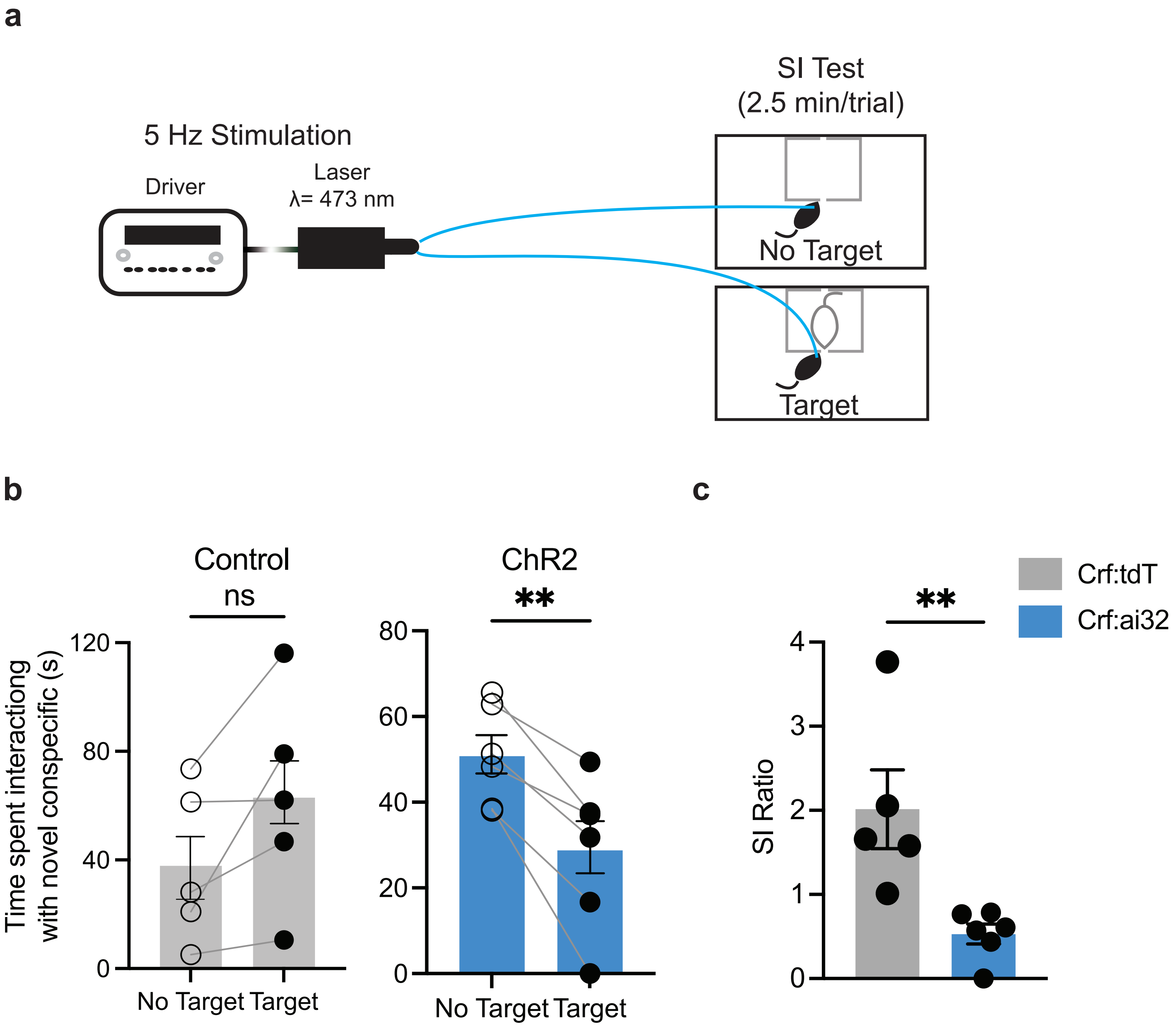


Acute BNSTov[CRF] stimulation induces social avoidance in mice with low stress exposure.

Extended Data Figure 5

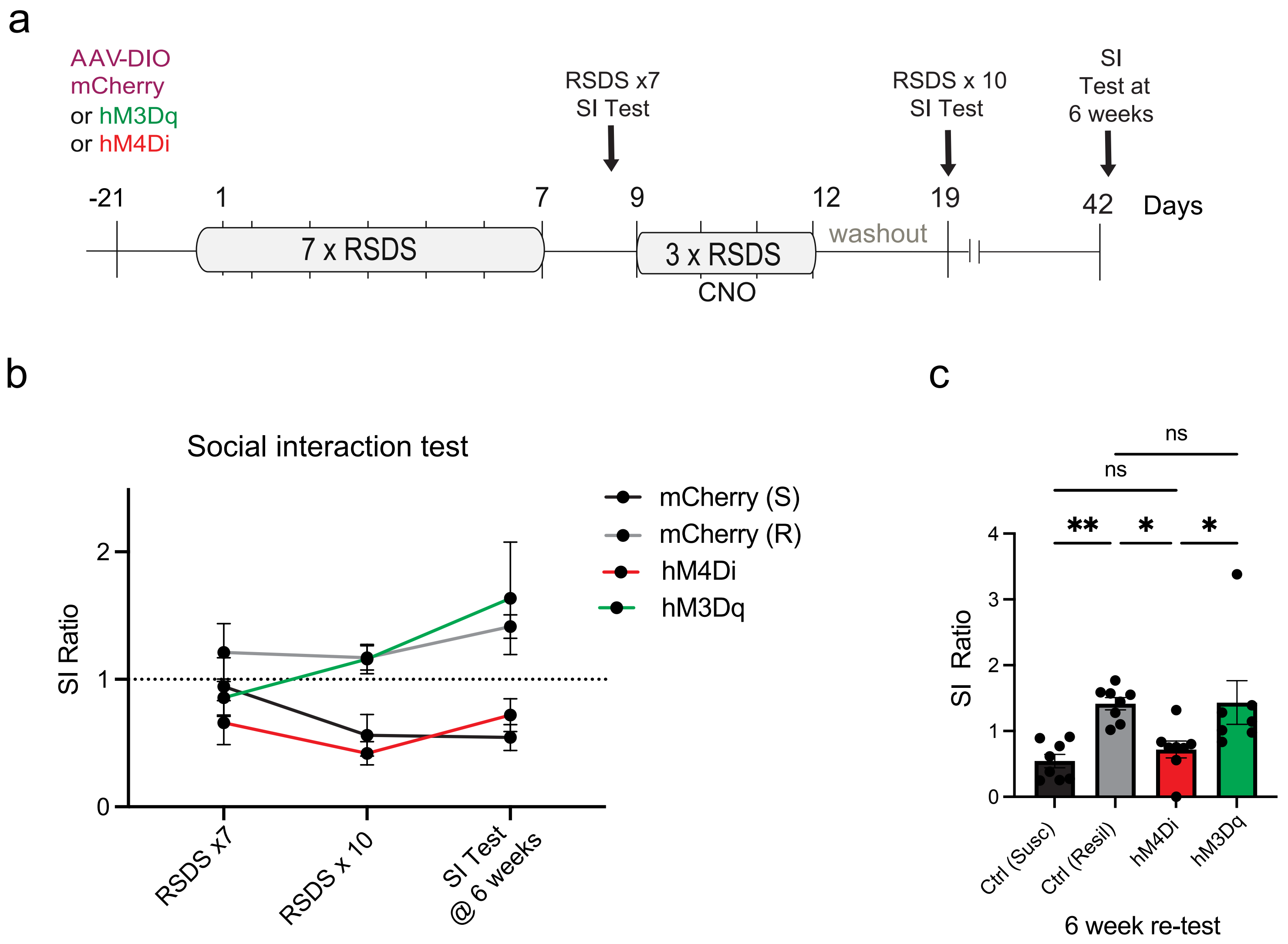


Bidirectional chemogenetic BNSTov$^{CRF}$ neuronal modulation produce durable susceptible /resilient phenotypes

Extended Data Figure 6

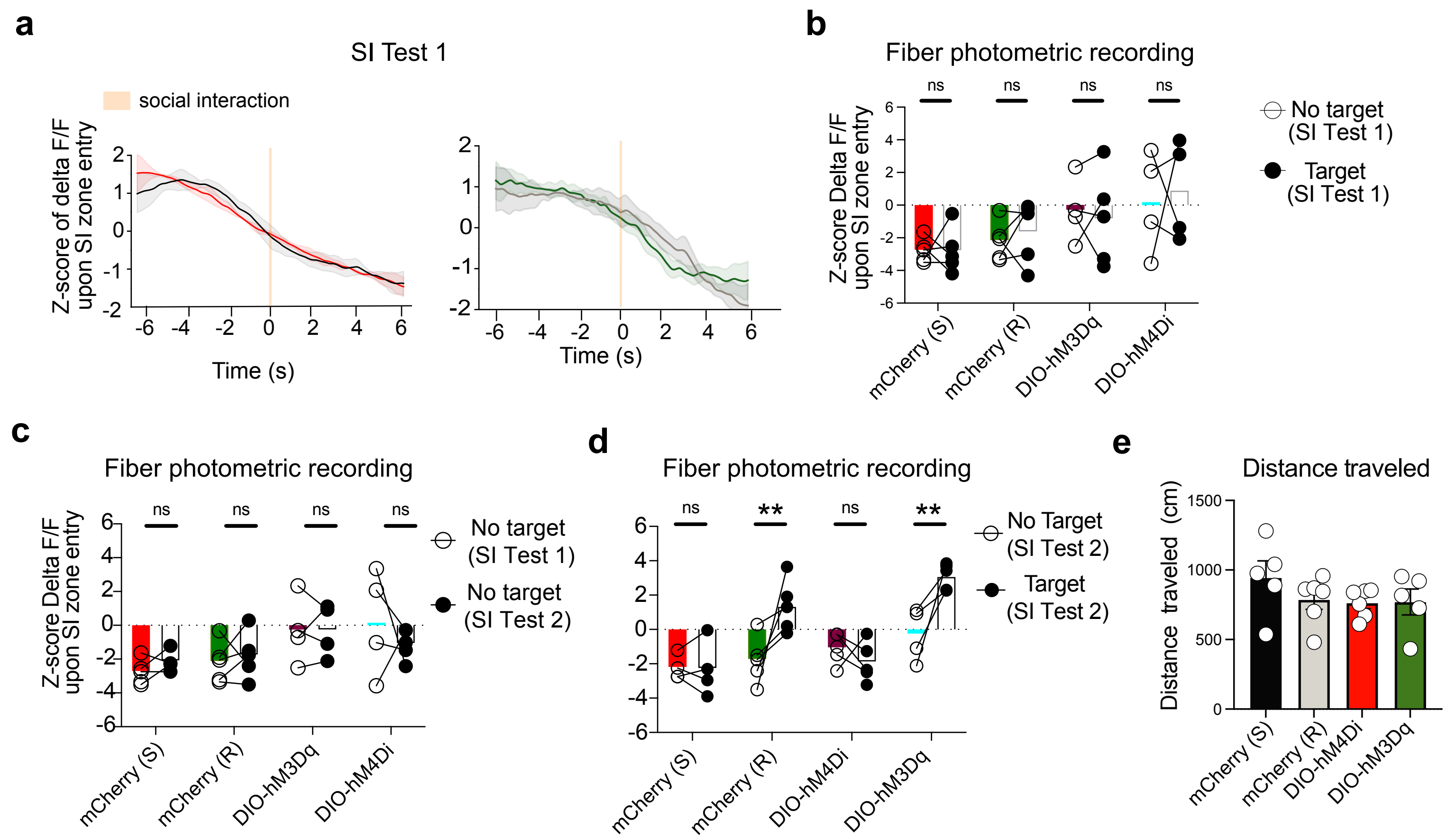


Prior to 10 SDEs, BNSTov CRF neurons are not activated in a novel social context

Extended data Figure 7

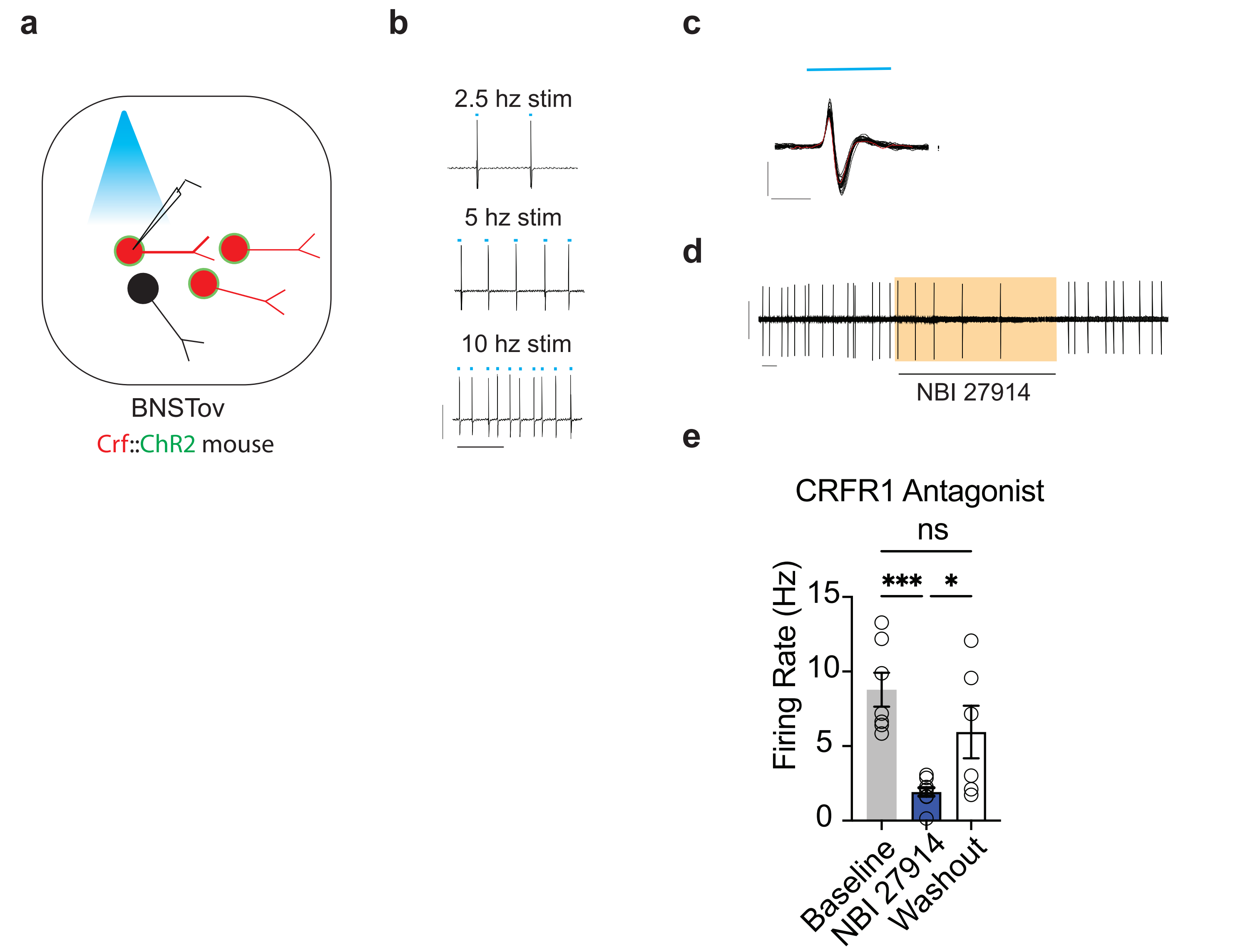


CRFR1 selective antagonist decreased spontanous firing rate in CRF+ neurons of optogenetically-induced resilient mice.

Extended Data Figure 8
