## Supplementary Figure Legends for "Stress History Modulates CRF Neurons to Establish Resilience"

**Supplemental Data Figure 1. Social defeat episodes, not days, promulgate the divergence of susceptible and resilient phenotypes.**

**A.** SI test of mice after 1, 4, 7, and 10 SDEs (n=5-18 mice/group). Two-way ANOVA, interaction $F_{(6,132)}$= 5.741, ****P<0.0001; row factor (episode #) $F_{(3,132)}$=12.10, ****P<0.0001; column factor (phenotype) $F_{(2, 132)}$=3.958, *P<0.05. Tukey's post-hoc test, Susceptible (1 vs 2 SDEs: *P<0.05; 1 vs 3 SDEs: ***P<0.001; 2 vs 4 SDEs: ****P<0.0001; 1 vs 4 SDEs: P=0.6295; 2 vs 3 SDEs: P=0.3994). Resilient (1 vs 2 SDEs: P=0.1114; 1 vs 3 SDEs: *P<0.05; 1 vs 4 SDEs: P=0.3072; 2 vs 3 SDEs: P=0.7298; 2 vs 4 SDEs: P=0.8999; 2 vs 4 SDEs: P=0.2929). Control (1 vs 2 SDEs: P=0.9899; 1 vs 3 SDEs: P=0.9999; 1 vs 4 SDEs: P=0.9998; 2 vs 3 SDEs: P=0.9954; 2 vs 4 SDEs: P=0.9966; 3 vs 4 SDEs: P=0.9999). **B.** Social interaction (SI) test of mice subjected to 7 episodes of social defeat stress (SDEs). Became susceptible vs became resilient, 2.062 ± 0.226 vs 1.967 ± 0.188; unpaired t-test, two-tailed t=0.3253, df=20, P=0.7483 (n=11 mice/group). **C.** Correlation of mice subjected to 10 SDEs (stressed x 10) and 7 SDEs (Stressed x7). $R^2$=0.06314, P=0.06314 (n=22 mice). **D.** Schematic of social defeat stress of 4 distinct cohorts as cross-sectional behavioral assessment of stress effect on SI. **E.** Social interaction test of cross-sectional behavioral assessment of 4 distinct cohorts of 1, 4, 7, 10 SDEs. Control=1.367 ± 0.09087 (n=28); SDE 1=0.9762 ± 0.05049 (n=10); SDE 4=1.377 ± 0.06854 (n=7); SDE 7=1.669 ± 0.1801 (n=16); SDE 10(R)=1.640 ± 0.1811 (n=13); SDE 10(S)=0.6164 ± 0.06406 (n=12). One-Way ANOVA $F_{(5,80)}$=8.341, ****P<0.0001. Sidak's post-hoc test, control vs 10: ***P<0.001; SDE 1 vs 7: *P<0.05; SDE 1 vs 10: *P<0.05; SDE 4 vs 10: *P<0.05; SDE 7 vs 10 (S): ****P<0.0001; SDE 10 (S) vs SDE 10 (R), ****P<0.0001. **F.** Sucrose preference test of independent cohorts of mice subjected to 1, 4, 7, or 10 SDEs. Control, 72.5160 ± 3.6657% (n=12); Stressed x7, 80.0383 ± 3.5376% (n=8); Susceptible, 51.7406 ± 5.716% (n=7); Resilient, 80.4516 ± 6.8360% (n=8). One-way ANOVA $F_{(3,31)}=6.337$, **P<0.01. Tukey's post-hoc test, control vs susceptible: *P<0.05; stressed x7 vs susceptible: **P<0.01; susceptible vs resilient: **P<0.01. **G.** Schematic of experimental design containing modified RSDS consisting of 10 episodes dispersed over 33 days, with intervention SI testing at days 9, 12, 30, and 35. **H.** 7 SDEs, SI Test 1: 1.966 ± 0.2245 (n=9); 7 SDEs, SI Test 2: 1.549 ± 0.179 (n=13); 7 SDEs, SI Test 3: 1.67 ± 0.1976 (n=16); 10 SDEs (S), SI Test 4: 0.5712 ± 0.121 (n=5); 10 SDEs (R), SI Test 4: 0.5712 ± 0.121 (n=8). One-Way ANOVA $F_{(4,46)}=4.320$, **P<0.01; Tukey's post-hoc test 7 SDEs/SI Test 2 vs 10 SDEs (S): **P<0.01; 7 SDEs/SI Test 2 vs 10 SDEs (S): *P<0.05; 7 SDEs/SI Test 3 vs 10 SDEs (S): *P<0.05. All data represent means ± SEM. *P<0.05, **P<0.01, ***P<0.001, ****P<0.0001, ns = not significant.

**Supplemental Data Figure 2: c-Fos immunohistochemistry of stress-sensitive regions reveals distinct patterns of activity to social interaction in the BNST.**
**A.** Experimental schematic, mice were subjected to 10 SDEs and a 15-minute non-physical social contact, 60 minutes after which brains were procured for c-Fos immunohistochemical analysis. **B.** Analysis of c-Fos immunoreactivity. Two-Way ANOVA brain interaction (region x phenotype) $F_{(10,50)}=2.247$, *P<0.05; brain region $F_{(5,50)}=0.7553$, P=0.5862; phenotype: $F_{(2,50)}=4.807$, *P<0.05; Tukey's post-hoc test:

BNST control vs resilient: *P<0.05; susceptible vs resilient: **P<0.01. **C.** Experimental schematic, mice were subjected to either 7 or 10 SDEs followed by a 15-minute non-physical social contact, 60 minutes after which brains were collected. **D.** Analysis of c-Fos immunoreactivity. Control: 45 ± 2.352 (n=6); stressed x7: 39.83 ± 3.911 (n=6); susceptible: 20 ± 5.206 (n=5); resilient: 84.4 ± 10.63 (n=5). One-Way ANOVA phenotype $F_{(3,18)}$=19.20, ****P<0.0001. Tukey's post-hoc test: control vs susceptible *P<0.05; control vs resilient: ***P<0.001; stressed x7 vs resilient: ***P<0.001; susceptible vs resilient: ****P<0.0001. Central Amygdala (CeA), Ventral Tegmental Area (VTA), Periaquactual Gray (PAG), Dorsal Raphe Nucleus (DRN), Lateral Hypothalamus (LH). All data represent means ± SEM. *P<0.05, **P<0.01, ***P<0.001, ****P<0.0001, ns = not significant.

**Supplemental Data Figure 3: Chemogenetic manipulation of $BNSTov^{CRF}$ neurons does not effect social defeat dynamics between aggressor CD-1 and C57/BL6J subordinate mice.**

**A.** Analysis of physical confrontational phases of the social defeat stress paradigm over SDEs 8-10. Two-Way ANOVA treatment x behavior $F_{(12,50)}$=0.2682, P=0.9917; treatment $F_{(3,50)}$=0.5199, P=0.6705; behavior $F_{(4,50)}$=28.4, ****P<0.0001 (n=14 mice). **B.** percentage of time spent exhibiting defensive behaviors associated with social defeat stress. mCherry (S): 44% cage exploration. 20% excessive grooming, 7% flight, 4% motionless, 25% fighting. mCherry (R): 47% cage exploration. 24% excessive grooming, 6% flight, 5% motionless, 18% fighting. hM4Di: 39% cage exploration, 27%

excessive grooming, 8% flight, 5% motionless, 21% fighting. hM3Dq: 44% cage exploration. 24% excessive grooming, 9% flight, 6% motionless, 17% fighting.

**Supplemental Data Figure 4: Chemogenetic manipulation of BNSTov$^{CRF}$ neurons bidirectionally modulate anxiety states.**

**A.** Experimental timeline. **B.** Representative heatmap of mice in EPM by treatment group. OA = open arm, CA = closed arm. **C.** Elevated plus maze analysis of time spent in open arm. mCherry (S), 0.2252 ± 0.22522 s (n=4); mCherry (R), 12.32 ± 7.1129 s (n=4); unpaired t-test, two-tailed, t=1.700, df=6, P=0.1401. hM3Dq, 42.1015 ± 14.3932 s (n=8); hM4Di, 1.95015 ± 1.7760 s (n=6); unpaired t-test, two-tailed, t=2.381, df=12, *P<0.05. **D.** Elevated plus maze analysis of number of entries. mCherry (S), 0.2500 ± 0.5000 (n=4); mCherry (R), 1.333 ± 2.3094 (n=3); unpaired t-test, two-tailed, t=0.9387, df=5, P=0.3910. hM3Dq, 12.125 ± 10.3845 (n=8); hM4Di, 0.6667 ± 1.2110 (n=6); unpaired t-test, two-tailed, t=2.662, df=12, *P<0.05. **E.** Experimental timeline, mice undergo RSDS with intervening SI tests and administered the open field test. SI test 1 and 2 depicted in gray denotes those findings are reported elsewhere in the report. **F.** Representative heatmap of open field behavior by treatment group. **G.** Time spent in the center of the open field arena. mCherry (S) vs mCherry (R), 10.35 ± 3.766 s (n=8), 5.926 ± 1.284 s (n=7); Mann Whitney test, two-tailed, P=0.9551. hM3Dq, 12.125 ± 10.3845 s (n=8); h4Di, 0.6667 ± 1.2110, *P<0.05 (n=7). All data represent means ± SEM. *P<0.05, ns = not significant.

**Supplemental Data Figure 5. Acute BNSTov$^{CRF}$ stimulation induces social avoidance in mice with low stress exposure.**

**A.** 5 Hz optogenetic 473 nm (blue light) photostimulation is induced with 2.5 minutes per trial with no target/target trials in tandem. **B.** Behavior of control mice during social interaction with novel conspecific within two (target vs no target) trials. Paired two-tailed t-test, t=2.282, df=4, P=0.0846 (n=5). ChR2 mice receiving photostimulatoin during social interaction test. Paired two-tailed t-test, t=5.389, df=5, **P<0.01 (n=6). **c**, SI ratio of control and ChR2 mice during the social interaction test. Crf:tdT and crf:ai32, unpaired two-tailed t-test, t=3.352, df=9, **P<0.01 (n=5-6). All data represent means ± SEM. *P<0.05, **P<0.01, ns = not significant.

**Supplemental Data Figure 6: Social interaction testing at 6 weeks following CNO modulation.**

**A.** Experimental timeline, mice are subjected to 7 and 10 SDEs, and administered an SI test after 7, 10, and six weeks from the start of the experiment. **B.** Display of animal behavior by treatment group over SI Testing. Dotting line at SI ratio = 1 demarcates the criteria for determining susceptible/resilient mice. SI ratio scores ≧ 1 defines resilient, <1 defines susceptible phenotypes. **C.** Social interaction test mCherry (S), 0.5439 ± 0.1019 (n=8); mCherry (R), 1.415 ± 0.09199 (n=8); hM4Di, 0.7200 ± 0.1286 (n=8); hM3Dq, 1.433 ± 0.3323 (n=7). One-Way ANOVA $F_{(3,27)}$=6.761, **P<0.01. Tukey's post hoc test mCherry (S) vs (R): **P<0.01; mCherry(S) vs hM3Dq: **P<0.01; mCherry(R) vs hM4Di: *P<0.05; hM4Di vs hM3Dq: *P<0.05. All data represent means ± SEM. *P<0.05, ns = not significant.

**Supplemental Data Figure 7: Prior to 10 SDEs BNSTov$^{CRF}$ neurons are not activated in a novel social context.**

**A.** Representative fiber photometric tracings of mice engaging with a novel conspecific during the social interaction test. **B.** Fiber photometric analysis of mice surrounding (-6 to +6) of an interaction bout. Two-Way ANOVA interaction (treatment x social target) $F_{(3,29)}$=0.14963, P=0.9291; treatment $F_{(3,29)}$=4.175, *P=0.0142; social target $F_{(1,29)}$=0.0611, P=0.8080 (n=4-5 mice/group). **C.** Fiber photometric analysis of recording during trials where social target was absent in both SI test 1 and 2. Two-Way ANOVA treatment x SI test $F_{(3,28)}$=0.6751, P=0.5746; treatment $F_{(3,28)}$=4.146, *P<0.05; social target $F_{(1,28)}$=0.01632, P=0.8993 (n=4-5 mice/group). **D.** Fiber photometric analysis of recording during trials when social target was absent vs present in SI Test 2. Two-Way ANOVA treatment (treatment x presence of social target) $F_{(3,28)}$=6.000, **P<0.01; treatment $F_{(3,28)}$ =13.44, ****P<0.0001; social target $F_{(1,28)}$=10.20, **P<0.01. Tukey's post-hoc test: mCherry (S) vs mCherry (R), *P<0.05; mCherry (S) vs hM3Dq, ****P<0.0001; hM4Di vs hM3Dq, ***P<0.001 (n=4-5 mice/group). **E.** Distance traveled. mCherry (S): 934.8633 ± 96.1767 cm (n=8); mCherry (R): 1278.478 ± 224.3500 cm (n=6); hM4Di: 1069.9941 ± 73.7286 cm (n=8). hM3Dq: 999.5635 ± 146.6165 cm (n=9). One-Way ANOVA treatment $F_{(3,16)}$=1.028, P=0.4066. All data represent means ± SEM. *P<0.05, **P<0.01, ***P< 0.001, ****P<0.0001, ns = not significant.

**Supplemental Data Figure 8. Optogenetic validation and CRFR1 selective antagonist decreased firing rate in CRF+ neurons of resilient mice. A.** Schematic of slice configuration of CRF cells in Crf::ChR2 mice. **B.** Optogenetic activation of varying firing frequency of 2.5 Hz, 5 Hz, 10 Hz firing fidelity, scale bar = 0.5 sec. **C.** Optogenetic induced waveform, scale bar (0.25 ms, 0.5 mV), blue bar symbolizes length of light pulse (0.7 ms, 5 mW). **D.** Sample trace of firing rate observed with bath application of NBI 27914 (CRFR1-specific antagonist). Scale bar is 1 sec, 0.5 mV. **E.** Firing rate of resilient mice (n= 3 mice). One-way ANOVA $F_{(2,19)}$=11.73, ***P<0.001. Tukey's post-hoc test: Baseline vs NBI 27914, ***P<0.001; NBI 27914 vs Washout *P<0.05. Baseline vs Washout P=0.1985. All data represent means ± SEM, *P<0.05, ***P<0.001, ns = not significant.
